# FAM210A mediates an inter-organelle crosstalk essential for protein synthesis and muscle growth in mouse

**DOI:** 10.1101/2023.08.03.551853

**Authors:** Jingjuan Chen, Feng Yue, Kun Ho Kim, Peipei Zhu, Jiamin Qiu, W. Andy Tao, Shihuan Kuang

## Abstract

Mitochondria are not only essential for energy production in eukaryocytes but also a key regulator of intracellular signaling. Here, we report an unappreciated role of mitochondria in regulating cytosolic protein translation in skeletal muscle cells (myofibers). We show that the expression of mitochondrial protein FAM210A (Family With Sequence Similarity 210 Member A) is positively associated with muscle mass in mice and humans. Muscle-specific *Myl1^Cre^*-driven *Fam210a* knockout (*Fam210a^MKO^*) in mice reduces mitochondrial density and function, leading to progressive muscle atrophy and premature death. Metabolomic and biochemical analyses reveal that *Fam210a^MKO^* reverses the oxidative TCA cycle towards the reductive direction, resulting in acetyl-CoA accumulation and hyperacetylation of cytosolic proteins. Specifically, hyperacetylation of several ribosomal proteins leads to disassembly of ribosomes and translational defects. Transplantation of *Fam210a*^MKO^ mitochondria into wildtype myoblasts is sufficient to elevate protein acetylation in recipient cells. These findings reveal a novel crosstalk between the mitochondrion and ribosome mediated by FAM210A.

## Introduction

Skeletal muscle growth and hypertrophy are contributed by the accretion of myonuclei from differentiated muscle stem cells (satellite cells) at early stages and by protein synthesis in the existing myofibers after the myonuclei addition ceases ^1, 2^. Skeletal muscle mass is maintained by a balanced action between protein synthesis and degradation. The most well studied pathway that leads to muscle mass gain is the mTOR pathway ^3^, where upon IGF-1 stimulation, Akt/mTOR/S6K/S6 are phosphorylated in a cascade that leads to the synthesis of rRNA/ribosomes and eventually protein synthesis. On the contrary, muscle protein degradation is controlled mainly by the ubiquitin-proteasome and autophagy-lysosome pathways. Understanding the mechanisms that regulate muscle growth provides potential strategies to boost muscle health for the improvement of life qualities.

Mitochondria are critical organelles supporting skeletal muscle metabolism ^4^. Several *in-vivo* genetic perturbation models highlight a key role of mitochondrial dynamics and function in skeletal muscle health. Proteins that control mitochondrial dynamics such as MFN2, OPA1, DRP1 regulate skeletal muscle mass and organismal health to a large extent ^5–8^. Similarly, mitochondrial biogenesis and transcriptional regulators such as PGC-1α and TFAM also govern muscle metabolic switches thereby affecting muscle mass ^9, 10^. Besides, mitochondrial metabolites provide intermediates for the biosynthesis of macromolecules ^4, 11, 12^, as well as ATP production for energy supply ^12^. Among the various metabolites, acetyl-CoA lies at the nexus of metabolic regulatory and signaling network ^13^. As the sole donor of protein acetylation, it affects muscle mass maintenance through post-translational modifications ^14, 15^. The crosstalk between mitochondria and other compartments such as nuclear ^16, 17^, lipid droplets ^18–21^ are being recognized as important events to maintain cellular homeostasis. Although intensive efforts have been taken to unravel the mechanisms linking mitochondrial fitness to muscle health, how mitochondrial metabolism regulates the postnatal muscle growth remains largely unclear.

The *Family with sequence similarity 210 member A* (*Fam210a*) gene was originally identified by genome-wide association studies to be correlated with bone density and muscle mass in children and adults ^22–24^. Paradoxically, *Fam210a* expression is very low in bone relative to that in the muscle tissue as shown by a *lacZ* reporter under the control of *Fam210a* promoter ^25^. Given the integral crosstalk between bone and muscle, and the abundant expression of *Fam210a* in the skeletal muscle, a more crucial function of FAM210A in the muscle is plausible. Indeed, whole-body heterozygous KO of *Fam210a* leads to reduced limb mass and muscle weakness, together with a deteriorated bone phenotype ^25^. Skeletal muscle-specific KO of *Fam210a* driven by inducible human alpha-skeletal actin (HSA)-Cre mice results in muscle weakness as well as a milder bone phenotype compared to the heterozygous KO, without affecting total body mass ^25^. Whether and how FAM210A regulates skeletal muscle growth and homeostasis remains elusive.

In the present study, using a skeletal muscle specific *Myl1* (myosin, light polypeptide 1) driven Cre recombinase specifically expressed in post-differentiation myocytes and multinucleated myofibers, we show that deletion of *Fam210a* caused a progressive myopathy and severe muscle weakness in mice, resulting in systemic metabolic defects and premature death. We find that loss of *Fam210a* disrupted the mitochondrial cristae structure and diminished the abundance of mitochondria in myofibers, accompanied by a deficiency in mitochondrial energy metabolism. Proteomics analysis reveals an induction of mitochondrial proteostastic response and apoptosis in *Fam210a*-null myofibers, concurrent with a reduction of the mitochondrial translation program. Further metabolomic analysis pointed to an abnormal flow of TCA cycle and accumulation of acetyl-CoA that leads to hyperacetylation of several ribosomal proteins which contributes to a stagnant translation in the *Fam210a*^MKO^ mice. These results uncover a novel role of FAM210A in regulating mitochondria metabolism and provide evidence linking metabolic inputs to the regulation of skeletal muscle growth and atrophy through protein acetylation.

## Results

### *Fam210a* is positively correlated with muscle mass in mice and humans

To verify the expression profile of FAM210A in various tissues, we performed western blot analysis using an antibody specifically recognizing FAM210A C terminus and found that FAM210A was highly expressed in tissues rich in mitochondria such as the heart, kidney, brown adipose tissue (BAT) and various skeletal muscles including diaphragm, tibialis anterior (TA), and soleus (**Figure S1A**). To determine how the expression of *Fam210a* is related to skeletal muscle mass, we analyzed publicly available microarray datasets on skeletal muscle biopsies from patients under various physiological and pathological conditions including aging, disuse, and muscular dystrophy, in which muscle atrophy occurs. Strikingly, compared to the young (20-29 years old) human subjects, *FAM210A* mRNA level in old (65-75 years old) was reduced by 37% and 37.5% in female and male muscle samples (GEO datasets GDS473 and GDS288), respectively (**Figure 1A**). In the vastus lateralis muscle of patients after 48 hours of knee immobilization (GEO dataset: GDS2083), *FAM210A* mRNA level was reduced by 33% compared to the control (**Figure 1B**). Similar downregulation of *FAM210A* mRNA was detected in patients with infantile-onset Pompe disease, a lysosomal storage disorder characterized by muscle weakness, as well as Duchenne Muscular Dystrophy (DMD) patients, an X-linked hereditary recessive myopathy (GEO datasets: GDS4410 and GDS611). Specifically, *FAM210A* mRNA level in muscles from Pompe disease and DMD was 18.6% and 42% lower than that in the healthy controls, respectively (**Figures 1C and 1D**).

**Figure 1.**
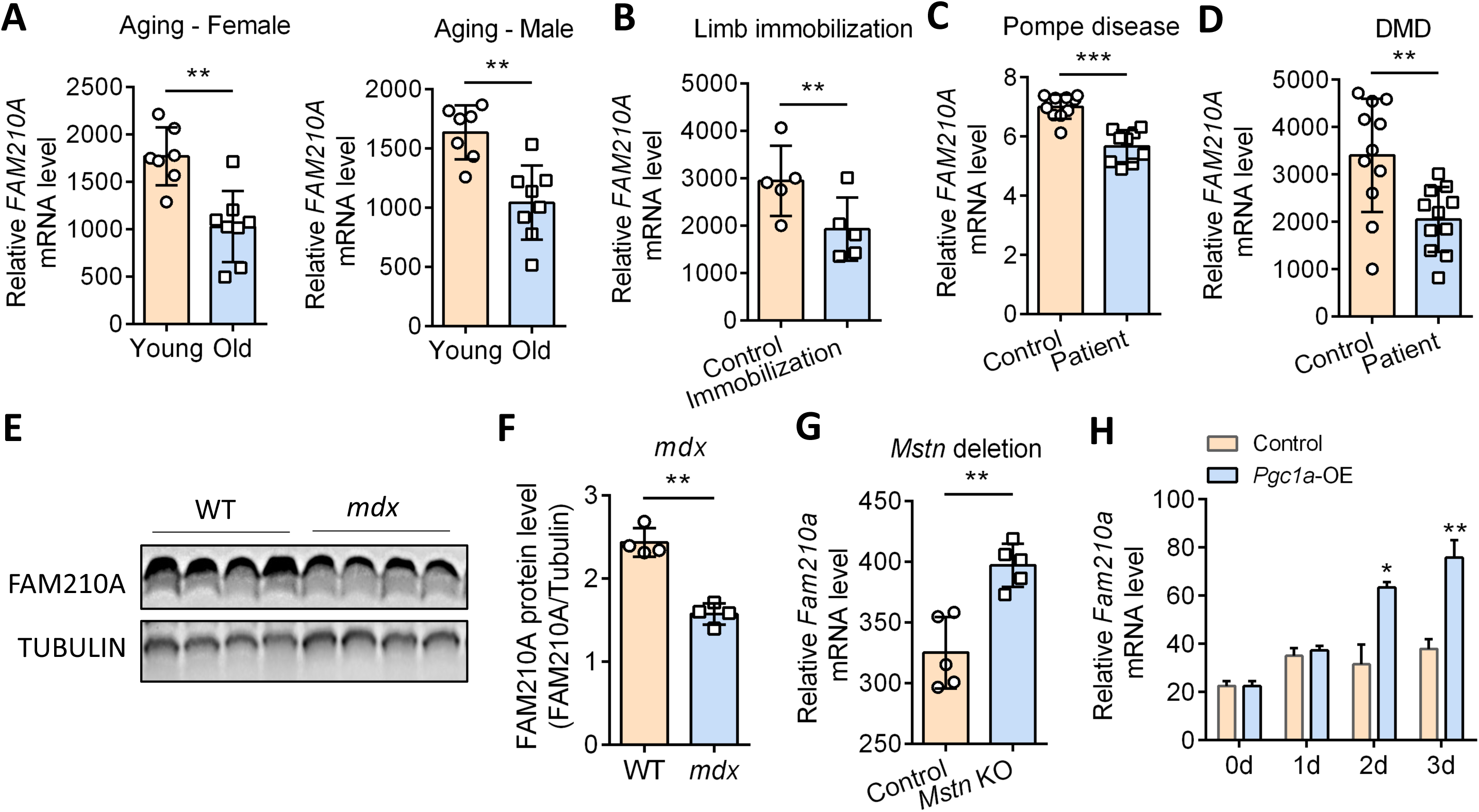
*Fam210a* expression decreases in atrophic and increases in hypertrophic skeletal muscles in patient and mouse. (**A**) mRNA level of *FAM210A* in young (20-29 year old) and old (65-75 year old) female and male human skeletal muscles. n = 7 for young, n = 8 for old. Datasets: GDS473 and GDS288. (**B**) mRNA level of *FAM210A* in vastus lateralis muscles of patients after knee immobilization for 48 hours. n = 5. Dataset: GDS2083. (**C**) mRNA level of *FAM210A* in infantile-onset Pompe patients with myopathy. n = 10 for control, n = 9 for Pompe patients. Dataset: GDS4410. (**D**) mRNA level of *FAM210A* in DMD patients. n = 11. Dataset: GDS61. (**E**) and (**F**) Western blot analysis and protein level of FAM210A in the soleus of D2.B10 *mdx* mice. n = 4. (**G**) mRNA level of *FAM210A* in the muscle of *myostatin* (*Mstn*) deletion mice. n = 5. Dataset: GDS3637. Student’s t-test, **p < 0.01. (**H**) Temporal mRNA level of *FAM210A* in the Pgc1a-overexpressed myoblasts during myogenic differentiation. n = 3 for 0d, n = 6 for 1-3d. Dataset: GDS1879. Student’s t-test, *p < 0.05, **p < 0.01, ***p < 0.001.

We further validated the changes of FAM210A protein in *mdx* mice, a model of human DMD. Consistent with the results in human patients, FAM210A protein level was 33% lower in the soleus muscle of *mdx* mice compared to the WT mice (**Figures 1E and 1F**). Conversely, an elevated expression of *Fam210a* mRNA was detected in hypertrophic muscles resulted from the *Myostatin* knockout (GEO dataset: GDS3637), with 21.5% higher than WT muscles (**Figure 1G**). Moreover, *Fam210a* mRNA expression was upregulated in newly differentiated myotubes that overexpressed the PPARG coactivator 1 alpha (PGC-1A) that drives myotube hypertrophy (**Figure 1H**, GEO dataset: GDS1879). Taken together, *Fam210a* level is positively corelated with muscle mass, as it is reduced in muscle atrophy conditions and increased in muscle hypertrophy conditions.

### Muscle-specific *Fam210a* deletion leads to muscle atrophy, growth retardation, and premature lethality

To understand the function of FAM210A in the skeletal muscles, we generated the *Myl1^Cre^*/*Fam210a^flox/flox^* (hereafter *Fam210a*^MKO^) mouse model by crossing *Fam210a^flox/flox^* mice with *Myl1^Cre^* mice to delete *Fam210a* specifically in muscle cells ^26, 27^. Specific and efficient deletion of *Fam210a* was validated by real-time PCR and western blot analyses (**Figures S1B and S1C**). The *Fam210a*^MKO^ mice were born at expected Mendelian ratios without apparent morphological and growth abnormalities before weaning. However, the *Fam210a*^MKO^ mice exhibited stagnant growth starting from the 4th week, with progressively smaller body size and hind limb muscles than those of the WT control mice (**Figures 2A and 2B**). Specifically, the *Fam210a*^MKO^ mice showed a 26.0% and 54.1% reduction in body weight (BW) compared to the WT controls at 4 and 6 weeks, respectively (**Figure 2B**). The growth retardation resulted in premature death at 8-9 weeks old (**Figure 2C**). Body composition analysis by EchoMRI detected that in comparison with the WT controls, the lean mass of the *Fam210a*^MKO^ mice showed a reduction at 32.5% and 56.6% at 4 and 6 weeks, respectively (**Figure 2D**), while only mild changes of the fat mass were observed (**Figure 2D**). However, the ratio of the lean mass and fat mass to body mass did not change between WT and *Fam210a*^MKO^ mice, indicating that the lean mass reduction was responsible for the body mass reduction (**Figure 2E**). Consistently, decreased TA muscle weight was observed in the *Fam210a*^MKO^ mice, with 42.5% and 75.4% reduction compared to the WT controls at 4 and 6 weeks, respectively (**Figure 2F**). These results indicate that FAM210A is required for the postnatal growth of skeletal muscle.

**Figure 2.**
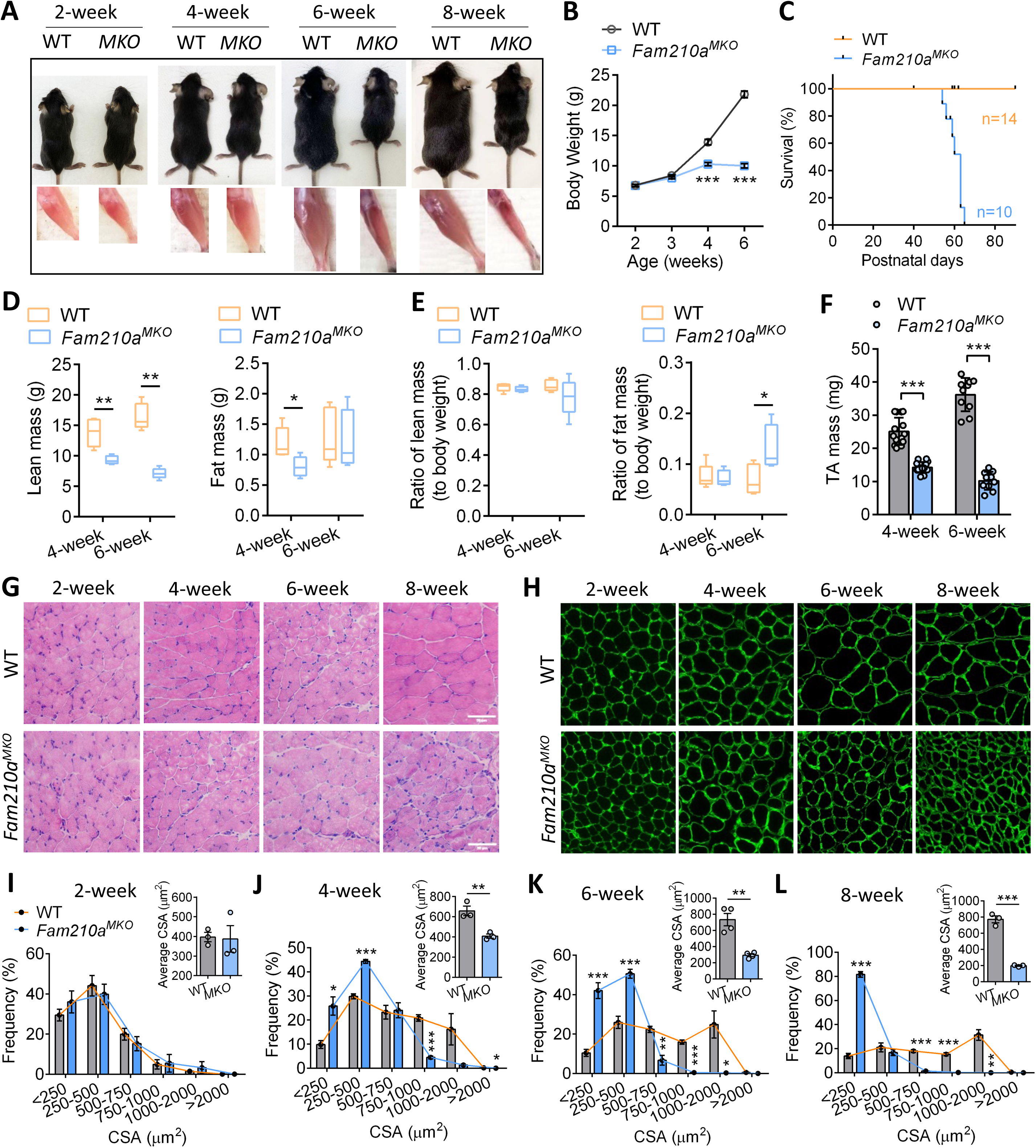
Myofiber-specific deletion of *Fam210a* impairs muscle growth, promotes muscle wasting and causes lethality. (**A**) WT and *Fam210a*^MKO^ mice and their respective hindlimbs at 2, 4, 6 and 8 weeks of age. (**B**) Body weight measurement of WT and *Fam210a*^MKO^ mice at different stages. (**C**) Survival curve of the WT and *Fam210a*^MKO^ mice. WT mice n= 14, Fam210a^MKO^ mice n=10. (**D**) EcoMRI scanning of lean mass and fat mass of the WT and *Fam210a*^MKO^ mice at 4 and 6 weeks of age. (**E**) Ratio of the lean mass and fat mass to body weight of the WT and *Fam210a*^MKO^ mice at 4 and 6 weeks of age. (**F**) TA muscle weight of the WT and *Fam210a*^MKO^ mice at 4 and 6 weeks of age. (**G**) H&E images of the muscle of the WT and *Fam210a*^MKO^ mice at different ages. (**H**) Laminin-staining of the TA muscle in WT and *Fam210a*^MKO^ mice at different ages. (**I-L**) Myofiber CSA distribution of the WT and *Fam210a*^MKO^ mice at different ages. (**I**) 2 weeks of age. (**J**) 4 weeks of age. (**K**) 6 weeks of age. (**L**) 8 weeks of age.

To determine the specific changes in the skeletal muscle tissues at the cellular level, we evaluated the histology in muscle cross-sections. Hematoxylin and eosin (H&E) staining did not reveal any obvious myofiber pathology in the *Fam210a*^MKO^ mice up to 8 weeks (**Figure 2G**). However, immunofluorescence analysis of basement membrane protein α-laminin that outline myofiber boundaries revealed fiber size abnormalities in the *Fam210a*^MKO^ mice (**Figure 2H**). Whereas the size of myofibers progressively increases from 2 to 8 weeks in WT mice, the myofibers stopped growing and then progressively decreased in size after 4 weeks in the *Fam210a*^MKO^ mice (**Figure 2H**). Unbiased quantification of myofiber cross sectional area (CSA) using ImageJ revealed that the average CSA was comparable in WT and *Fam210a*^MKO^ mice at 2 weeks old (**Figure 2I**), but the average CSA of *Fam210a*^MKO^ mice became progressively (40% - 75%) smaller than that of WT mice from 4 to 8 weeks (**Figures 2I-2L**). Strikingly, the average CSA of the *Fam210a*^MKO^ mice reduced from 387.1 μm^2^ to 197.0 μm^2^ within 6 weeks from 2-week-old to 8-week-old (**Figures 2I-2L**), indicating progressive muscle wasting. Consistently, the CSA distribution patterns were indistinguishable in WT and *Fam210a*^MKO^ at 2 weeks but the portion of larger myofiber in *Fam210a*^MKO^ mice was significantly lower than that of WT starting at 4-week-old (**Figure 2J**). At 6 and 8 weeks, the differences between CSA distribution curves of WT and *Fam210a*^MKO^ mice were more prominent: while >60% of WT myofibers were larger than 500 μm^2^, only <10% of *Fam210a*^MKO^ myofibers reached this size (**Figures 2K and 2L**). Together, these results demonstrate severe growth retardation and progressive muscle atrophy in the *Fam210a*^MKO^ mice.

### Loss of *Fam210a* causes severe muscle weakness

We next investigated how *Fam210a*^MKO^ affects muscle function. As severe muscle wasting inevitably impairs contractile function, we examined WT and *Fam210a*^MKO^ mice at the onset of the muscle atrophy (4 weeks old), when the mutant mice only showed a slight reduction of body weight and muscle mass (**Figure 2**). When the mice were subjected to endurance training on treadmill, the maximum running speed, total running time, and total running distance of *Fam210a*^MKO^ mice were 33.3%, 46.8% and 61.2% lower than those of sex-matched WT littermates, respectively (**Figures 3A-3C**). We further subjected the mice to grip strength measurement and found the *Fam210a*^MKO^ mice also manifested a 55% reduction of grip strength (0.64 N) compared to their WT littermates (1.41 N) (**Figure 3D**). To determine whether the deficiencies of motor function and grip strength in *Fam210a*^MKO^ mice are specifically due to muscle dysfunction, we assessed the contractility of the extensor digitorum longus (EDL, fast-twitch) and soleus (SOL, slow-twitch) muscles. In response to increasing frequencies of electric stimulation, the absolute contractile force significantly increased and plateaued in the WT mice, but the rate of increase and maximal forces of the *Fam210a*^MKO^ EDL and SOL muscles were significantly lower (**Figures 3E and 3G**). Specifically, the maximal absolute force of EDL and SOL showed a 68.9% and 32.8% reduction in *Fam210a*^MKO^ mice (**Figures 3F** and **3H**). Consistent results were observed in specific contractile force when the absolute contractile force was normalized by the muscle CSA (**Figures 3I and 3K**), where the maximal specific force of EDL and SOL manifested a decrease at 42.1% and 19.4% in *Fam210a*^MKO^ mice (**Figures 3J** and **3L**). Taken together, these results provide compelling evidence that the loss of FAM210A in muscle leads to severe defect in muscle contractile function.

**Figure 3.**
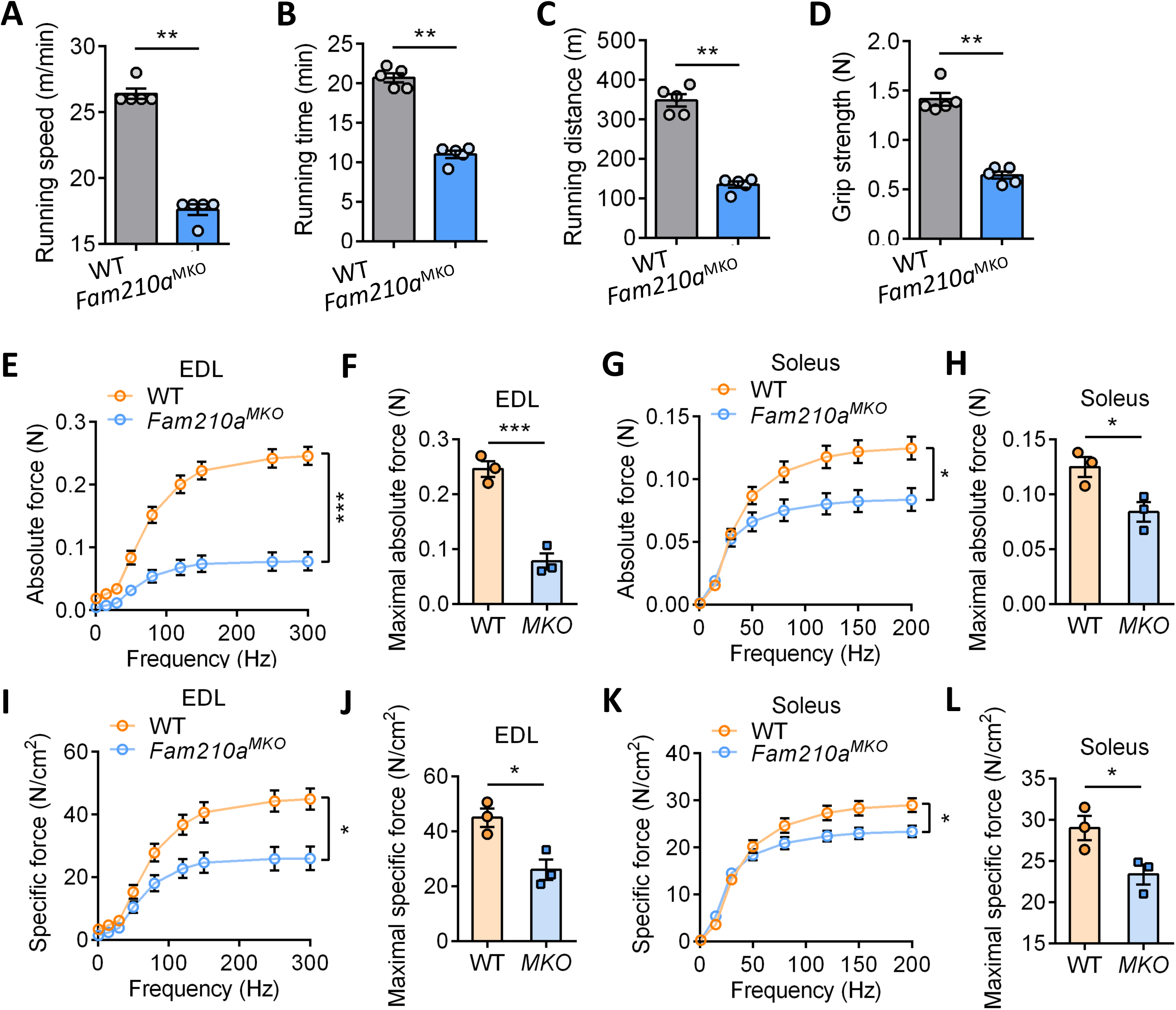
*Fam210a* deletion causes muscle weakness. (**A**) Treadmill running speed of the WT and *Fam210a*^MKO^ mice at 4 weeks of age. N=5. (**B**) Treadmill running time of the WT and *Fam210a*^MKO^ mice at 4 weeks of age. N=5. (**C**) Treadmill running distance of the WT and *Fam210a*^MKO^ mice at 4 weeks of age. N=5. (**D**) Grip strength of the WT and *Fam210a*^MKO^ mice at 4 weeks of age. N=5. (**E**) Muscle contraction measurement of absolute force in EDL muscle in WT and *Fam210a*^MKO^ mice. (**F**) Quantification of the maximal absolute force in EDL, related to (**E**). (**G**) Muscle contraction measurement of absolute force in SOL muscle in WT and *Fam210a*^MKO^ mice. (**H**) Quantification of the maximal absolute force in SOL, related to (**G**). (**I)** Muscle contraction measurement of specific force in EDL muscle in WT and *Fam210a*^MKO^ mice. (**J**) Quantification of the maximal specific force in EDL, related to (**I**). (**K**) Muscle contraction measurement of specific force in SOL muscle in WT and *Fam210a*^MKO^ mice. (**L**) Quantification of the maximal specific force in SOL, related to (**K**).

### *Fam210a*^MKO^ mice exhibit systemic metabolic defects

Besides motor function, skeletal muscle contributes substantially to metabolic health as muscle accounts for ∼40% of total body weight and ∼70% of total protein and absorbs ∼75% of post prandial glucose ^11, 28^. We therefore monitored the respiratory activity of the mice using the indirect calorimetry. The *Fam210a*^MKO^ mice showed a significantly higher levels of lean mass-normalized oxygen consumption (VO_2_) and carbon dioxide production (VCO_2_) during both day and night times (**Figures S2A and S2C**). Quantitatively, the average VO_2_ of the *Fam210a*^MKO^ mice was 19.4% and 27.6% higher than the WT control while the average VCO_2_ was 19.0% and 26.1% higher in the *Fam210a*^MKO^ mice in day and night times, respectively (**Figures S2B and S2D**). Moreover, the heat production of the *Fam210a*^MKO^ mice was significantly lower during the daytime but only showed a tendency to decrease in the nighttime (**Figure S2E**). Because muscle serves as sink for glucose regulation ^11^, we monitored the blood glucose level and clearance capacity of these mice by glucose tolerance test (GTT). The fasting glucose level was significantly lower in *Fam210a*^MKO^ mice (47.3 mg/dL) compared to the WT mice (123 mg/dL) (**Figure S2F**). Despite a much slower glucose clearance rate, the glucose tolerance of the *Fam210a*^MKO^ mice was slightly better than that of the WT mice, likely due to the 65% lower fasting glucose level (**Figures S2G and S2H**). These results suggest that the loss of FAM210A in skeletal muscle causes systemic metabolic dysfunction.

### *Fam210a*^MKO^ increases oxidative myofibers and decreases glycolytic myofibers

Metabolic properties (glycolytic or oxidative) of myofibers are intimately related to exercise performance and systemic metabolism. We next investigated the myofiber compositions in EDL and SOL using myosin heavy chain (MyHC) isoform specific monoclonal antibodies (**Figure S3A**). In mice, the oxidative capacity declines while glycolytic capacity increases gradually in type I, IIa, IIx/d, IIb MyHC. We observed a gradual increase in the portion of Type IIa fiber and the hybrid Type I/IIa fibers in the EDL and SOL muscles of the *Fam210a*^MKO^ mice from 4 weeks to 8 weeks, respectively (**Figures S3A and S3B**). This was correlated to a decrease in the glycolytic IIx myofibers in the SOL muscles (**Figure S3B**). We further quantified the myofiber fiber sizes of different fiber types and found that the decrease of fiber size was consistent across all fiber types in SOL muscle (**Figure S3D**). However, the slowest fiber type (Type IIa) did not atrophy compared to other faster fiber types in EDL muscle (**Figure S3C**). These results suggest that *Fam210a*^MKO^ induces a metabolic switch to a more oxidative state.

We then directly assessed the mitochondrial oxidative capacity by staining succinate dehydrogenase (SDH) activity in the TA muscle. Surprisingly, we observed a progressive decrease of the SDH enzymatic activity from 2 to 8 weeks in the *Fam210a*^MKO^ mice (**Figure 4A**). Specifically, the SDH activity staining signal was comparable between WT and *Fam210a*^MKO^ mice at 2 weeks old, however it was remarkably lower in *Fam210a*^MKO^ muscles at 4, 6 and 8 weeks as can be visualized by the less and lighter blue staining compared to the WT muscles (**Figure 4A**). The reduction of SDH activity corroborated with reduced expression of OXPHOS proteins, notably the SDHB protein (**Figure 4B**).

**Figure 4.**
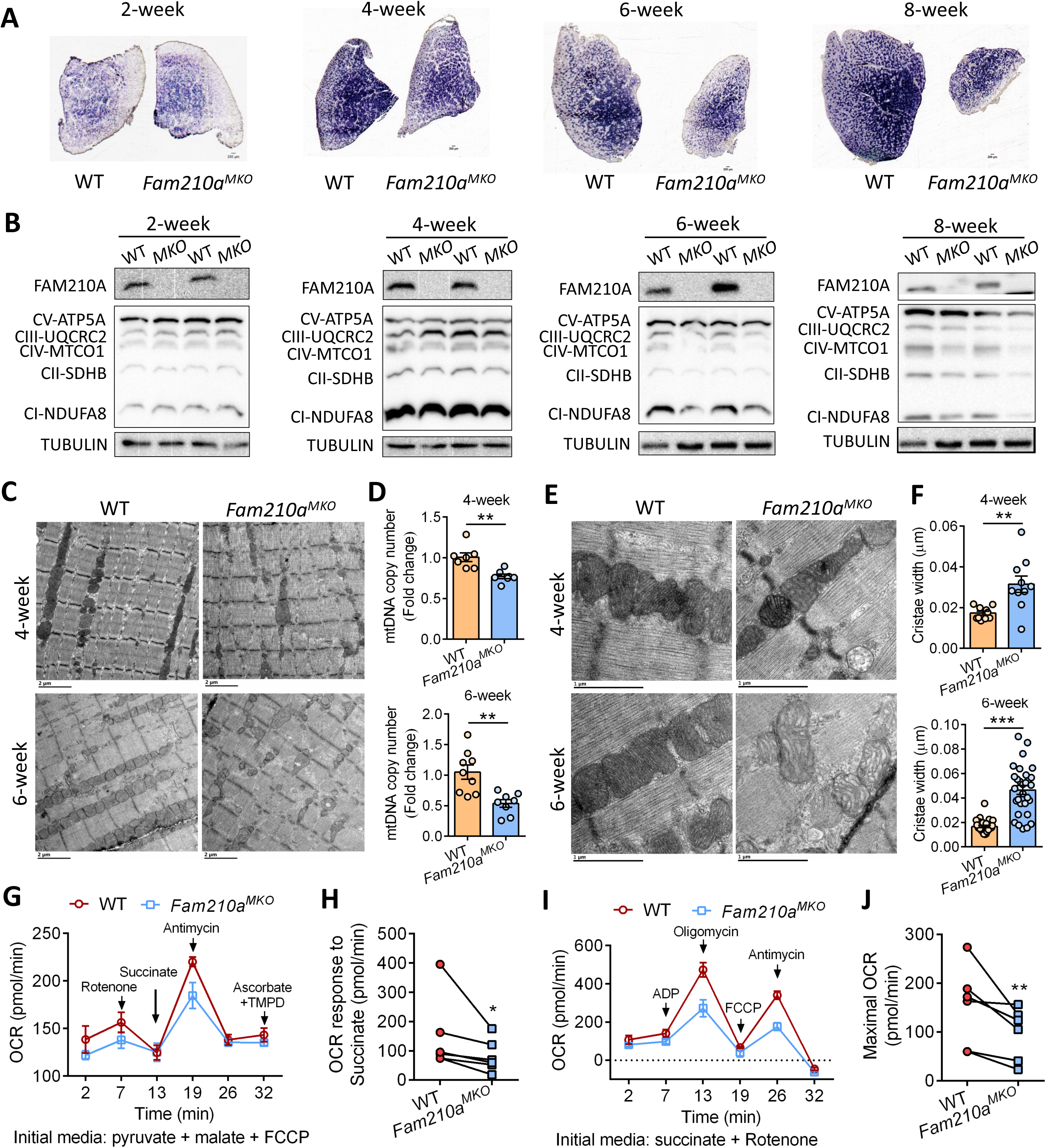
*Fam210a* deletion results in mitochondrial morphological and energetic defects. (**A**) Succinate dehydrogenase activity staining (SDH) in the TA muscle at different ages in WT and *Fam210a*^MKO^ mice. (**B**) Muscle mitochondrial protein analysis by western blot in WT and *Fam210a*^MKO^ mice at different ages. (**C**) Representative TEM images of the muscle showing mitochondria abundance in the WT and *Fam210a*^MKO^ mice at 4 and 6 weeks of age. (**D**) mtDNA quantification by qPCR at 4 and 6 weeks of age in WT and *Fam210a*^MKO^ mice. (**E**) Representative TEM images of the mitochondria morphology in WT and *Fam210a*^MKO^ mice at 4 and 6 weeks of age. (**F**) Cristae distance measurement in the WT and *Fam210a*^MKO^ mitochondria at 4 and 6 weeks of age from TEM images. (**G**) Representative graph of electron flow experiment using purified mitochondria from WT and *Fam210a*^MKO^ muscle, drug administrations were indicated in the graph. (**H**) Quantification of the oxygen consumption rate after the administration of succinate, related to (**G**). (**I**) Representative graph of coupling experiment using purified mitochondria from WT and *Fam210a*^MKO^ muscle, drug administrations were indicated in the graph. (**J**) Quantification of the oxygen consumption rate after the administration of FCCP, related to (**I**).

Additionally, the level of mitochondrial encoded respiratory chain protein MTCO1 was also lower progressively in the *Fam210a*^MKO^ mice **(Figure 4B**). Taken together, these results indicate that *Fam210a*^MKO^ muscles undergo metabolic adaptation that uncouples muscle oxidative/glycolytic properties from MyHC subtypes.

### *Fam210a*^MKO^ disrupts mitochondrial cristae structures and causes respiratory dysfunction

To directly explore how the loss of FAM210A affects mitochondria, we examined the mitochondrial abundance in *Fam210a*^MKO^ mice. Quantification of mitochondrial DNA (mtDNA) by qPCR revealed a gradual reduction of mtDNA in the *Fam210a*^MKO^ mice, with a 23.5% and 48.9% reduction at 4-weeks and 6-weeks respectively (**Figure 4D**). To further visualize the morphological changes of mitochondria, we performed transmission electron microscope (TEM) imaging. Consistent with the reduction in mtDNA copy numbers, *Fam210a*^MKO^ muscle contained fewer and smaller mitochondria per field at 4-week-old (**Figure 4C**). Morphologically, whereas a packed lamellar cristae structure was observed in the WT mitochondria, the *Fam210a*^MKO^ mitochondria contained irregular tubular-vesicular cristae with increased cristae distance, defined as the largest distance between two adjacent cristae (**Figure 4E**). A more severe mitochondrial abnormality was observed in *Fam210a*^MKO^ mice at 6-week-old, manifested by the higher occurrence of irregular tubular-vesicular cristae, indicating the progressive defects of mitochondria. Specifically, the cristae distance in the mitochondria increased 0.58 and 1.37-fold at 4-week and 6-week-old muscle, respectively (**Figure 4F**).

To address whether the *Fam210a*^MKO^ mitochondria with the structural alterations were energetically defective, we conducted an electron flow experiment with isolated mitochondria using Seahorse Analyzer, where the function of each individual complex was interrogated sequentially ^29^. In the presence of the pyruvate, malate and uncoupler FCCP, no significant changes of the OCR for basal respiration and Complex I activity (after the addition of Rotenone) were observed. Notably, the OCR stimulated by the addition of succinate was significantly decreased in *Fam210a*^MKO^ mitochondria, suggesting that the SDH-mediated Complex II function is defective (**Figures 4G and 4H**). Moreover, we examined the energy coupling by stimulating the mitochondria with the substrate ADP and FCCP. We didn’t observe a reduction in ADP-stimulated respiration, but the *Fam210a*^MKO^ mitochondria had reduced maximal respiration upon the addition of FCCP (**Figures 4I and 4J**). These results implicate FAM210A in the maintenance of mitochondria integrity and energy production.

### *Fam210a*^MKO^ induces a proteome featuring mitochondrial apoptotic response and impaired mitochondrial respiration and translation program

To gain a mechanistic insight into how FAM210A regulates mitochondrial cristae integrity and energy metabolism, we performed proteomic analysis using mitochondria isolated from WT and *Fam210a*^MKO^ muscles at 6 weeks of age. Principal Component Analysis (PCA) showed a tight clustering of the WT and *Fam210a*^MKO^ samples (**Figure 5A**). A total of 161 proteins were differentially expressed between *Fam210a*^MKO^ and WT mitochondria (p-Value < 0.05; Fold Change > 1.5). Among the 53 proteins upregulated in the *Fam210a*^MKO^ mitochondria, an induction of apoptotic proteins such as BAX, diablo IAP-binding mitochondria protein (DIABLO), and growth hormone inducible transmembrane protein (GHITM) ^30^ were observed (**Figure 5B**). These results suggest that the *Fam210a*^MKO^ mitochondria may trigger retrograde signaling to the nucleus to induce apoptotic response to remove the damaged mitochondria ^31^. On the other hand, among the 108 decreased proteins, we observed a significant reduction of the mitochondrial translation machinery TUFM and several mitochondrial ribosomal proteins (**Figure 5B**).

**Figure 5.**
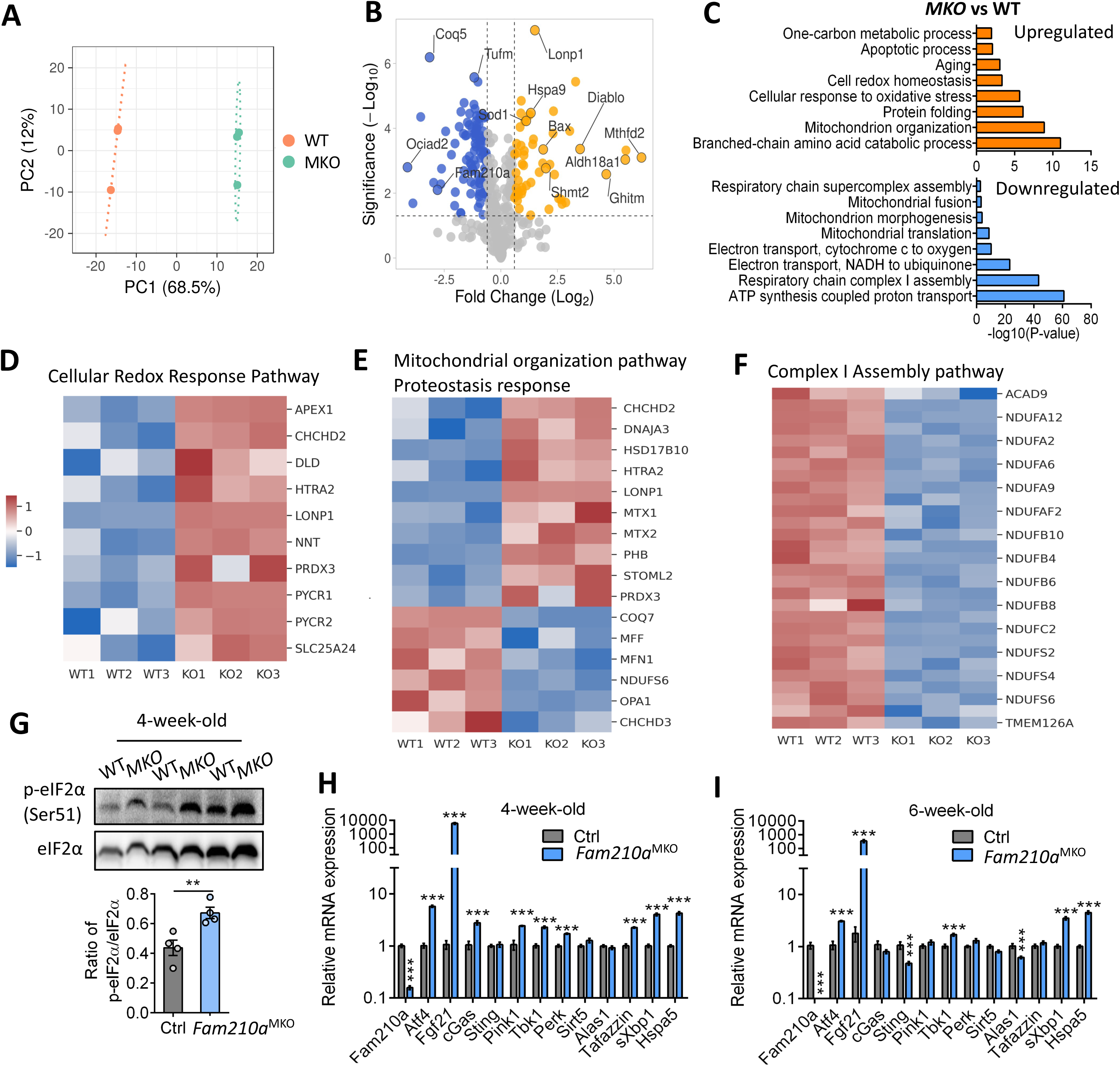
*Fam210a* deletion leads to proteomic stress and elicits ISR signaling. (**A**) PCA plot of the proteomics data from 6 weeks old mitochondria isolated from WT and *Fam210a*^MKO^ muscle. (**B**) Volcano plot of the differentially expressed proteins in purified mitochondria from WT and *Fam210a*^MKO^ mice muscle at 6 weeks of age. (**C**) Pathway analysis of the down-regulated and up-regulated biological pathways in the *Fam210a*^MKO^ mitochondria. (**D-F**) Heatmaps of the differentially expressed proteins featuring cellular redox response pathway (**D**), mitochondrial organization/proteostasis pathway (**E**) and complex I assembly pathway (**F**). (**G**) Western blot analysis of the ISR pathway effector p-eIF2α in 4-week-old muscle from WT and *Fam210a*^MKO^. Quantification provided in the bar graph. (**H**) q-PCR analysis of the ISR pathway from 4-week-old muscle from WT and *Fam210a*^MKO^ mice. (**I**) q-PCR analysis of the ISR pathway from 6-week-old muscle from WT and *Fam210a*^MKO^ mice.

To unravel the function of the differentially expressed proteins, we performed Gene Ontology (GO) pathway analysis. The top downregulated biological pathways enriched in the *Fam210a^MKO^* mitochondria include ATP synthesis-coupled proton transport and Complex I assembly (**Figure 5C**), which corroborated with previous western blots showing reduced Complex I protein (**Figure 4B**) and Seahorse energetic results showing compromised mitochondrial function (**Figures 4I and 4J**). Interestingly, we found enrichment of the biological pathways related to redox response, fatty acid oxidation (FAO), mitochondrial organization and protein folding in the *Fam210a^MKO^* mitochondria (**Figures 5D-5F**). Given the observed deficiency of the TCA cycle and the electron transport chain, the increase of FAO pathway, which serves to utilize cellular lipids, might be a compensatory response. The up-regulated mitochondrial organization may also indicate an adaptive response to the abnormal mitochondrial cristae structure (**Figure 4E**).

As the integrated stress response (ISR) pathway is well studied to be induced under various mitochondrial stress ^32, 33^, we examined the protein and mRNA level of the effector components of the ISR pathway. The phosphorylated eukaryotic initiation factor 2 α (phospho-eIF2αSer 51) level was increased in the *Fam210a*^MKO^ compared to the WT (**Figure 5G**). Consistently, the mRNA levels of *Atf4*, *Fgf21* and the associated transcripts were highly induced in the *Fam210a*^MKO^ both in the 4-week and 6-week-old muscle samples (**Figures 5H and 5I**). Collectively, proteomics and biochemical analysis reveal that FAM210A is essential in maintaining the proteostasis in the mitochondria.

### *Fam210a*^MKO^ disrupts the flow of TCA cycle, causing abnormal accumulation of acetyl-CoA

Mitochondrial TCA cycle and electron transport chain (ETC) are interconnected by the succinate dehydrogenase enzyme complex located at the mitochondrial inner membrane (**Figure 6A**). With the previous Seahorse energetic results and SDH staining results pointing to the metabolic defects centered on TCA cycle, we conducted a targeted metabolomics analysis of TCA cycle intermediates. Intriguingly, we detected a 200% higher α-ketoglutarate level (**Figure 6B**) but a 40% lower succinate level (**Figure 6C**) in the *Fam210a*^MKO^ muscle compared to the WT control, indicating that there was a blockage in the conversion between α-ketoglutarate to succinate (**Figure 6A**). Indeed, to identify FAM210A interacting proteins related to TCA cycle and complex II activity, we found by IP-MS/MS analysis (**Figure S4A**) that SUCLG2, the β-subunit in the succinyl-CoA synthetase complex, was among the top hits (**Figure S4B**). We further investigated the enzymatic activity of SUCL complex and detected an increase of SUCL enzymatic activity (**Figure 6D**), which catalyze the bidirectional conversion between succinate and succinyl-CoA (**Figure 6A**). Furthermore, the blockade in the TCA cycle led to the buildup of acetyl-CoA in the *Fam210a*^MKO^ muscle as shown by metabolomics data (**Figure 6E**). Together, these results show that FAM210A is critical in maintaining the TCA cycle metabolic balance.

**Figure 6.**
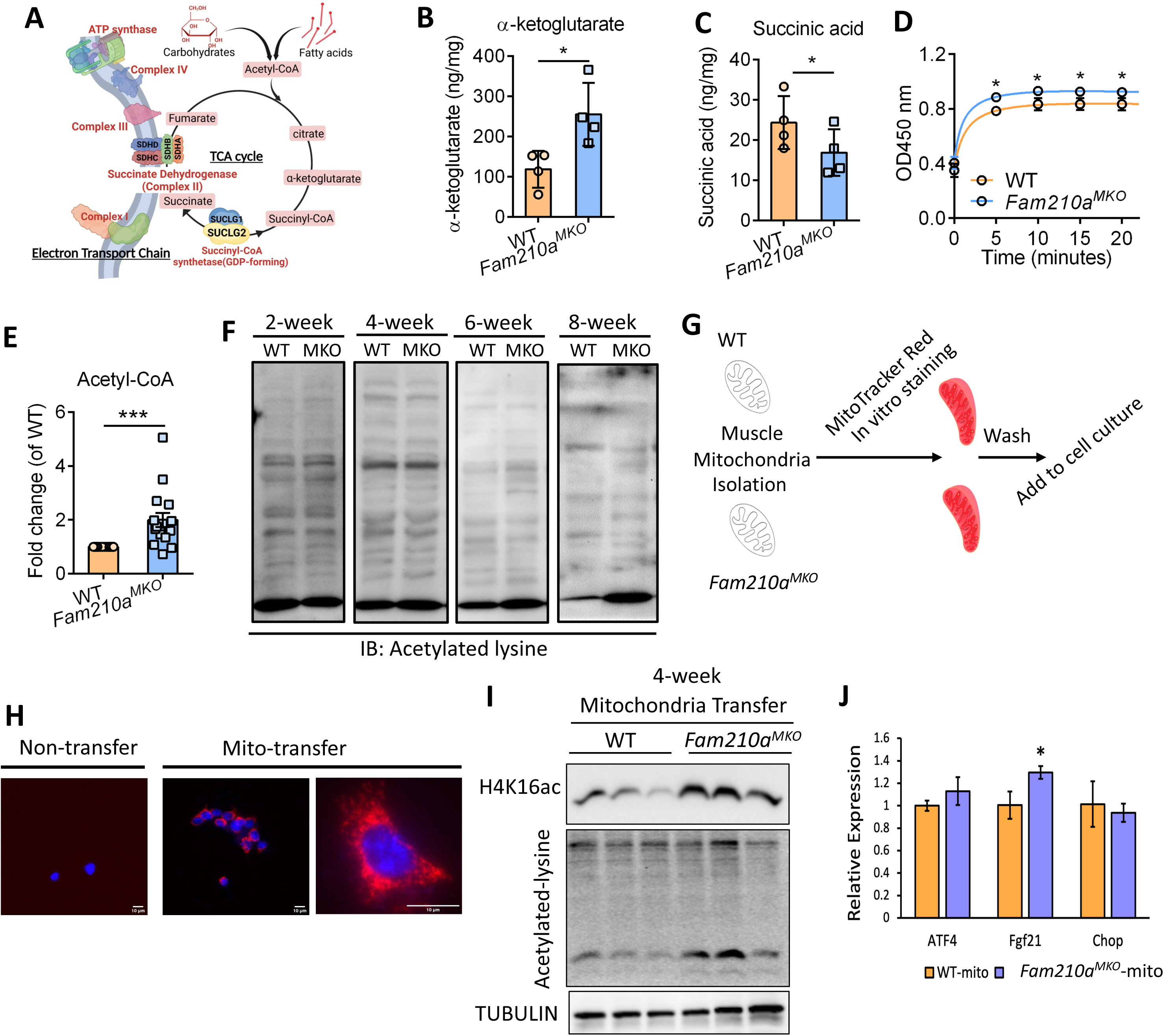
*Fam210a* deletion in mitochondria leads to metabolic defects and induce protein hyperacetylation. (**A**) Diagram of the intersection between TCA cycle and Electron transport chain in the mitochondria. (**B**) Targeted metabolomics analysis of α-ketoglutarate level in the muscle lysate at 4 weeks of age in WT and *Fam210a*^MKO^ mouse. (**C**) Targeted metabolomics analysis of succinate in the muscle lysate at 4 weeks of age in WT and *Fam210a*^MKO^ mouse. (**D**) Succinyl-CoA ligase enzymatic activity measurement in muscle lysate at 4 weeks of age in WT and *Fam210a*^MKO^ mouse, OD450nm representing the relative enzymatic activity. (**E**) Acetyl-CoA abundance quantification in the WT and *Fam210a*^MKO^ muscle at 4 weeks of age by targeted metabolomics analysis. (**F**) Protein acetylation profile analysis using pan-acetyl-Lysine antibody in muscle lysates in different ages in WT and *Fam210a*^MKO^ mice. (**G**) Experimental design of the mitochondrial transfer experiment. (**H**) MitoTracker Red staining after mitochondria transplantation for 24hours in WT recipient myoblasts. (**I**) Western blot analysis of the acetyl-Lysine profile in the WT recipient myoblast protein lysate after mitochondria-transfer from WT and *Fam210a*^MKO^ muscle. (**J**) q-PCR analysis of the WT recipient cells after being transplanted with WT or *Fam210a*^MKO^ mitochondria.

### FAM210A-null mitochondria induce protein hyperacetylation in WT cells

Given acetyl-CoA is a major donor for protein acetylation, and the significant increase of acetyl-CoA pool in *Fam210a*^MKO^ muscle, we then assessed the acetylome of the proteins in the *Fam210a*^MKO^ muscle. Western blot analysis using anti-acetylated lysine antibody showed no significant protein acetylation change between WT and *Fam210a*^MKO^ at 2- and 4-weeks, however a progressive increase of acetylation of several proteins especially at the 15 kDa size range was observed in 6- and 8-weeks (**Figure 6F**). To test if the increase of acetylation in the *Fam210a*^MKO^ muscle was directly driven by mitochondrial defects, we performed transplantation assay to transfer MitoTracker Red labeled WT or *Fam210a*^MKO^ mitochondria into the WT primary myoblasts (**Figure 6G**). When 0-16 μg of isolated mitochondria were incubated with 30,000 recipient cells for 24 hour, MitoTracker Red signal revealed a dose-dependent transfer of labeled mitochondria into recipient cells, starting at 2 μg (**Figure 6H**). The MitoTracker Red fluorescence signals coincided with the relative levels of mtDNA in the cells as quantified by q-PCR (Figure S5A**, blue bar**). Moreover, the mitochondria copy number increase elicited by the transfer of extrinsic mitochondria was to a lesser extent at 48 h post transplantation compared to 24 h (Figure S5A**, orange bar**). Based on the results, we transferred 8 μg mitochondria per 30,000 recipient cells and measured uptake dynamics, which revealed efficient mitochondria transfer in 30 min (Figure S5B). We then evaluated the acetylation profile of the recipient primary myoblasts and observed that the protein acetylation at the 15 kDa size was consistently increased (**Figure 6I**). To rule out the possibility that the elevated acetylation reflects proteins carried by the transferred mitochondria into the donor cells, we blotted the total lysate with nuclear H4K16ac of the host origin. Consistently, we saw an increase of H4K16ac level in the primary myoblasts transplanted with *Fam210a*^MKO^ mitochondria (**Figure 6I**). When we examined the cellular response in the recipient cells, we observed that the transcript level of the ISR signaling pathway component *Fgf21* was significantly increased and *Atf4* was marginally increased by transfer of *Fam210a*^MKO^ mitochondria (**Figure 6J**). These results provide compelling evidence that FAM210a*-*null mitochondria are sufficient to drive aberrant acetylation of cytosolic and nuclear proteins in normal cells.

### *Fam210a* deletion leads to hyper-acetylation of cytosolic ribosomal proteins

The specific increase of protein acetylation at the 15 kDa size prompted us to perform IP-MS/MS to identify the proteins that are hyper-acetylated in the *Fam210a*^MKO^. Acetylated proteins were pulled down using antibody against the acetylated lysine residue and separated by SDS-PAGE gel (**Figure 7A**). The proteins in the gel at 10-20 kDa were excised and subjected to MS/MS identification (**Figure 7B**). This led to the identification of several histone proteins and ribosomal proteins that were hyper-acetylated in the *Fam210a*^MKO^ muscles (**Figure 7C**). We further purified the ribosomes from muscle tissues of WT and *Fam210a*^MKO^ and blotted them with anti-acetylated lysine antibody to check the level of acetylation in ribosomal proteins. We observed that the ribosomal proteins were increasingly hyper-acetylated with age in the *Fam210a*^MKO^ compared to the WT ribosomes (**Figure 7D**).

**Figure 7.**
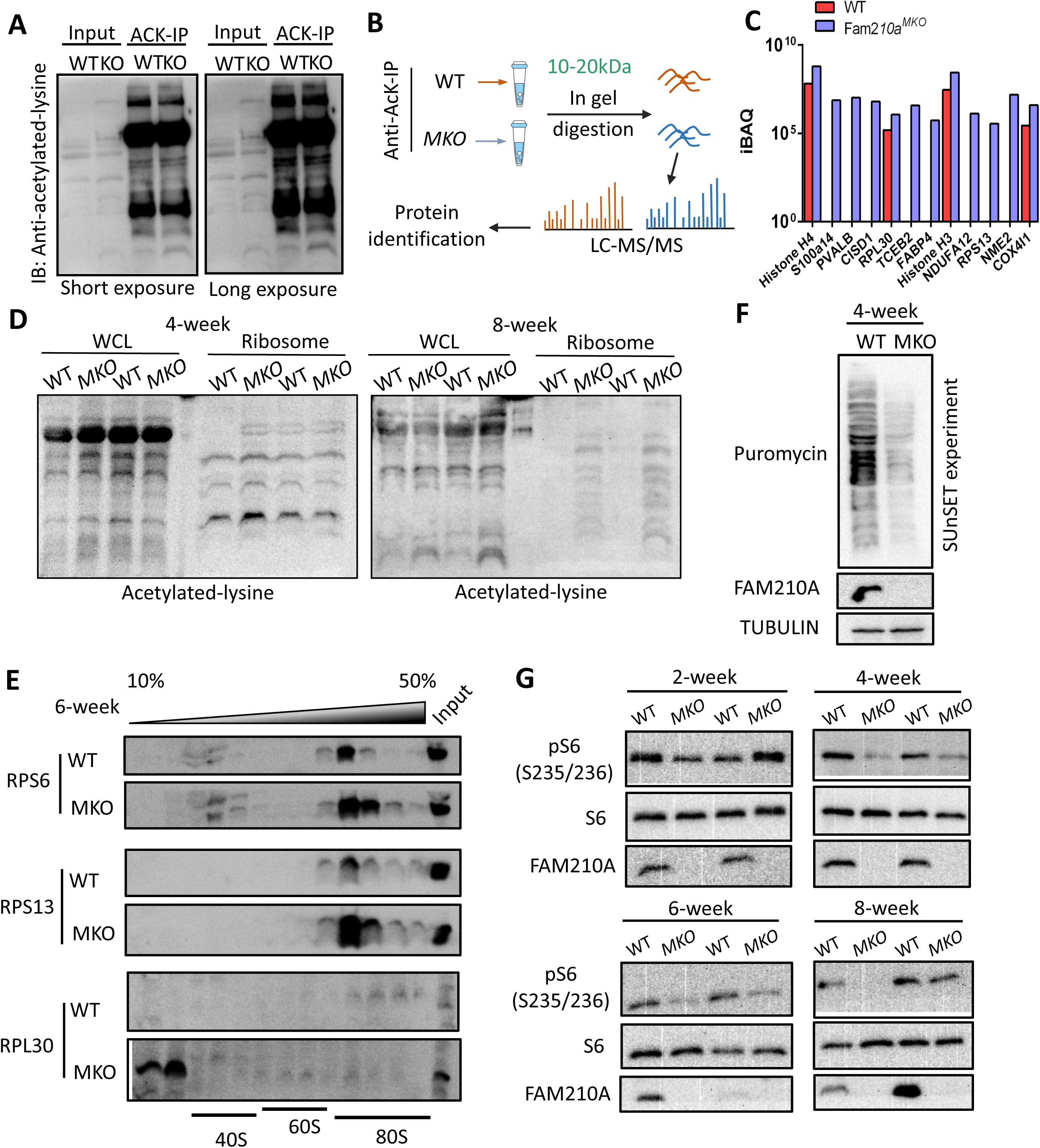
*Fam210a* deletion leads to ribosomal protein hyperacetylation and protein translation defects. (**A**) Acetyl-Lysine antibody immunoprecipitation to enrich the hyper-acetylated proteins in *Fam210a*^MKO^ muscle. (**B**) Workflow of the LC-MS/MS to identify the 15KDa range proteins that were highly acetylated in the *Fam210a*^MKO^ muscle compared to WT. (**C**) Identification of the ribosomal proteins and histone proteins from the 15kDa proteins shown in (**A**). (**D**) Ribosome isolation from muscle and verification of the acetyl-lysine protein in the ribosomal fraction at 4 and 8 weeks of age. (**E**) Ribosome profiling assaying of the assembly of ribosomal proteins into the different subunits in the muscle ribosomes at 6 weeks of age. RPS6 and RPS13 were used as markers for the ribosomal small subunits and RPL30 were used as markers for the ribosomal large subunits. (**F**) SUnSET experiment revealing the incorporation of puromycin in the muscle in WT and *Fam210a*^MKO^ muscle at 4 weeks of age. (**G**) Western blot analysis showing the reduction of phospho-S6 in muscle lysate in different ages in WT and *Fam210a*^MKO^ mice.

To investigate the functional consequence of the hyperacetylation of ribosomal proteins, we performed ribosomal profiling assay to examine the assembly. While it appeared that the small subunit markers such as RPS6 and RPS13 were more abundant in the 40S and polysome factions in the *Fam210a*^MKO^ samples, large subunit marker RPL30 was retained in the top fractions as “free” proteins after the gradient centrifuge (**Figure 7E**). The *Fam210a*^MKO^ samples also exhibited a relative increase of RPL30 in the 60S faction rather than the polysome fraction compared to the WT muscles (**Figure 7E**). Because ribosomes are translational machineries, we performed SUnSET assay ^34, 35^, which labels nascent proteins with puromycin to monitor the translation efficiency in the muscles. This assay revealed reduced levels of puromycin-labeled nascent polypeptide chains in the *Fam210a*^MKO^ muscle (**Figure 7F**). Consistently, we observed a reduction of phospho-S6 level progressively in *Fam210a*^MKO^ muscle (**Figure 7G**), indicative of reduced translation. Since the ISR gene signature was also elevated in the *Fam210a*^MKO^ muscle (**Figures 5G-5I**), it could further dampen the translation defects. These results demonstrate that *Fam210a* deletion in skeletal muscle induces the hyper-acetylation of several small ribosomal proteins that leads to ribosomal disassembly and translational deficiency.

## Discussion

Mitochondria metabolism is important for ensuring that the skeletal muscles meet the energetic demands for its motor function and retro-signaling to the nucleus for organismal homeostasis. Here in the study, we demonstrate that FAM210A is indispensable for postnatal muscle growth and maintenance. Loss of *Fam210a* leads to deregulation of the mitochondrial cristae organization and further proteostatic stress. FAM210A deficiency in the mitochondria disrupts the normal TCA cycle which shifts the equilibrium of the oxidative reaction towards the reductive fashion. This leads to the accumulation of acetyl-CoA and hyperacetylation of ribosomal proteins in the cytosol and histone proteins in the nucleus. Hyperacetylation of ribosomal proteins disrupts the ribosomal assembly and protein translation, eventually muscle atrophy ensues. These results pinpoint the significance of metabolic control on the function of ribosomes and cytosolic translation through mitochondrion and ribosome crosstalk.

A previous study reported that global deletion of *Fam210a* leads to premature death in utero at E9.5 days prior to myogenesis, highlighting its critical role in early development ^25^. While tamoxifen inducible *Fam210a* global KO at postnatal 28 days leads to substantially decreased both lean mass and body weight, skeletal muscle-specific KO induced at postnatal 28 days under the HSA promoter only mildly affects the lean mass but not the body weight ^25^. Nonetheless, in our *Myl1* driven *Fam210a* deletion model, we observed a drastic reduction of both lean mass and body weight in the *Fam210a*^MKO^ mice starting at 4 weeks of age, which provided compelling evidence on the importance of *Fam210a* in muscle homeostasis. The milder phenotype seen in the HSA-Cre driven *Fam210a* deletion might be due to the efficiency of the exogenous promoter as well as the timing of the induction of the KO, compared to the *Myl1* gene whose expression is readily detectable in skeletal muscle at E12.5 and almost 96% of myonuclei expressed *Myl1* at postnatal day 12 ^36^. The mild phenotype before 4 weeks of age might be due to the non-essential role of mitochondrial metabolism during embryonic and perinatal myogenesis. Indeed, several reports have revealed a critical role of glycolysis on promoting cell proliferation and muscle size in *Drosophila* during embryonic myogenesis ^37–40^. The reliance on aerobic glycolysis rather than oxidative respiration during developmental stage can explain the inconsistency between the expression profile of *Myl1* and the onset of the pathology observed in our model. In addition, nuclei-accretion mediated by satellite cells underlies the hyperplasic growth of myofibers until 3 weeks of age ^2^, but *Myl1*^Cre^ mediated *Fam210a*^MKO^ only affect post-differentiating cells without affecting satellite cells. These results demonstrate that FAM210A plays a critical role in maintaining muscle mass and function.

Moreover, we are the first to characterize how FAM210A regulate mitochondria structural and proteostatic homeostasis. However, the detailed mechanisms underlying the aberrant mitochondrial structure remain unclear. Some plausible factors could be energetic stress and the metabolic imbalance. Loss of membrane potential, overproduction of ROS, cytochrome c and mtDNA release to the cytosol all can induce signaling cascades disrupting mitochondrial quality control ^17, 41^. A previous study has identified *Fam210a* as the target of miR-574 and showed that FAM210A is important in maintaining the cardiac mitochondrial translation program. In the cardiac model, FAM210A down-regulation leads to the reduction of membrane potential and ATP production. In our finding, FAM210A controls the metabolic balance by regulating the Succinyl-CoA Synthetase (SCS) complex activity and promotes the oxidative flow of the TCA cycle. Therefore, *Fam210a* KO in skeletal muscle leads to the unmet energetic demands and accumulation of the upstream metabolite acetyl-CoA and subsequently elicit the mitochondrial quality control program. However, the detailed mechanism through which FAM210A regulates SCS enzyme complex remains to be investigated.

In our finding, FAM210A mediated mitochondrial metabolism controls the level of acetyl-CoA and the absence of such regulation leads to overproduction of acetyl-CoA which fail to be oxidized. We reasoned that the energetic deficiency experienced by the myofibers will signals through the ISR pathway and induce the translational arrests by phosphorylating eIF2α ^32, 33, 42^. Moreover, when the proportion of ribosomal protein phosphorylation buildup in the *Fam210a*^MKO^, the assembly of the translational machinery is impaired therefore further dampen the cellular translation. Interestingly, another study has implicated FAM210A in the regulation of mitochondrial-encoded protein synthesis in cardiac muscle through its interaction with ATPase Family AAA Domain Containing 3A (ATAD3A) and mitochondrial Elongation Factor Tu (EF-Tu) ^43^. In the *Fam210a*^MKO^ model, we observed a decrease of EF-Tu protein in the proteomics data, as well as a decreased MTCO1 protein level by western blot, which also pinpoint a regulation of the mitochondrial translation machinery upon FAM210A loss in skeletal muscle. These results suggest that FAM210A can modulate mitochondrial translation in addition to regulating the cytosolic translation machinery through a mitochondrion-ribosome crosstalk. The difference between skeletal muscle and cardiac muscle on FAM210A’s working mechanism might be due to metabolic preference of the tissues, which requires further investigation to see if these two pathways diverge from a common upstream effector or that the inherent mitochondrial metabolic difference govern the different mode of action.

Acetyl-CoA serves as the messenger that links the cellular metabolic state to the cellular processes ^44–46^. Mounting evidence have suggested a direct link between acetyl-CoA production and global protein acetylation. Interestingly, ribosomal protein acetylation has been charactered in organisms including bacteria, yeast, and rice ^47–52^. Xu *et al*. using point-mutagenesis have shown that mimicking acetylation led to decreased ribosomal protein stability while depleting acetylation by substitution to arginine residues on the ribosomal proteins led to increased stability in rice ribosomes ^53^. In *Fam210a*^MKO^ model, FAM210A regulates the SCS enzymatic activity and the outcome is the compensational overproduction of the acetyl-CoA, thereby increasing the protein acetylation and hampering the ribosome assembly. Together with ISR activation and ribosomal assembly defects, the translation halts in the *Fam210a*^MKO^ myofibers, and the muscle mass decreases due to the net loss of protein. We have identified a new regulatory axis which links metabolic input to ribosomal homeostasis and muscle mass maintenance. The ribosomal protein acetylation sites of the target proteins, however, needs to be further dissected to have a clear understanding of such regulation.

In summary, we have revealed the critical role of FAM210A in the maintenance of mitochondrial structural regulation and metabolic balance by mediating a mitochondrion-ribosomal crosstalk, which contributes to skeletal muscle mass maintenance and functions.

## Materials and Methods

### Experimental Design

### Animals

The *Fam210a*^flox/flox^ mouse was generated by Nanjing Biomedical Research Institute of Nanjing University (NBRI) in a C57BL/6J background. The *Myl1*^Cre^ mouse was generously provided by Steven Burden (Skirball Institute of Biomolecular Medicine, NYU). *Fam210a*^flox/flox^ mice were bred to *Myl1*^Cre^ mice to generate a muscle specific knockout mouse model. Mice were housed in an animal facility with free access to water and standard rodent chow diet. All procedures involving mice were approved by the Purdue Animal Care and Use Committee (PACUC). Male or female mice were used and always gender matched for each specific experiment.

### Whole-Body Composition Analysis

The fat and lean mass compositions of mice were analyzed with an EchoMRI 130 analyzer (EchoMRI, Houston, TX, USA). Briefly, the instrument was first calibrated using corn oil. Total body fat, lean mass, free water and total water were measured in a non-invasive manner. The measurement tubes were disinfected with 70% ethanol between each mouse.

### Treadmill, Grip Strength and Indirect calorimetric measurement

A detailed protocol for treadmill and grip strength tests of mice was published elsewhere^54^. The endurance test was performed using and Exer-3/6 treadmill (Columbus Instruments) as previously described. Briefly, the mice were trained for 3 consecutive days before the test at 10 min per day at 10 m/min, +10 slope, and 3 Hz + 1mA electrical stimulation. For the test, each mouse was placed in a dedicated lane and started to run on the treadmill at 10 m/min for 5 min, with a gradual increase of speed at 2 m/min every 2 min until the mice were exhausted. Exhaustion was determined when the mouse was unable to run on the treadmill for 10 s despite mechanical prodding. The maximum speed, total running distance and running time achieved were recorded by Treadmill Software (Columbus Instruments).

The grip strength meter (Columbus Instruments) was used to measure four-limb grip strength in mice following the manufacturer’s guide. Briefly, mice were acclimated for 10 min before the test in the hood. The mouse was allowed to grab onto the metal pull bar with the paws. Then the mouse tail was gently pulled backward until mice could not hold onto the bar. The force at the time of release was recorded as the peak tension. Each mouse was tested 7 times. The grip strength was defined as the average strength.

For indirect calorimetric measurement, mice were acclimated to the system (Oxymax, Columbus Instruments) for at least 24 hours before measurement. The system was kept in a stable environment at 22°C with a 12-h light cycle (6 am to 6 pm). Mice were placed in individual chambers with free access to food and water. The data were presented as corrected energy expenditure levels. Average energy expenditure of day (6 am to 6 pm) and night (6 pm to 6 am) were calculated as the mean values of all data points in the time period.

### Muscle Contractile Force Measurement

Muscle contractile force measurement was conducted by using an *in-vitro* muscle test system (1200A intact muscle test system, Aurora Scientific) ^55^. In brief, the mice were anesthetized with isoflurane, and the hind limb was cut and immediately placed in a bicarbonate-buffered solution (137 mM NaCl, 5 mM KCl, 1 mM MgSO_4_, 1 mM NaH_2_PO_4_, 24 mM NaHCO_3_, and 2 mM CaCl_2_) equilibrated with 95% O_2_-5% CO_2_ (pH 7.4) for dissection. The EDL muscle and the SOL muscle were dissected with care to ensure the tendons were intact. After isolation, the proximal and distal tendons were tied with braided silk suture thread (4-0, Fine Science Tools). The muscle was then mounted in the muscle test system continuously bubbled with carbogen (5% CO_2_ in O_2_) at room temperature. The stimulation protocol consisted of supramaximal electrical current delivered through platinum electrodes using a biphasic high-power stimulator (701C, Aurora Scientific). After determining optimal length (Lo) with a series of twitch stimulations at supramaximal voltage, the temperature of the organ bath was increased to 32°C. After 10 min of thermal equilibration, a force-frequency relationship was generated using selected frequencies between 1 and 300 Hz for the EDL muscle (200 ms train) and 1 and 200 Hz for the soleus muscle (500 ms train). This was followed by a 5 min fatiguing protocol of 500 ms volleys of 40 Hz stimulation applied once every 2 s. The muscle was then removed from the organ bath, trimmed of connective tissue, blotted dry, and weighed. Muscle CSA was determined by dividing the wet muscle mass by the product of Lo and muscle-specific density (1.056 g/cm^3^). Specific force (N/cm^2^) was calculated by dividing the muscle force (N) by the CSA (cm^2^). The maximum twitch response was analyzed for peak twitch tension, time to peak twitch tension, and twitch half-relaxation time.

### Transmission Electron Microscopy

Isolated extensor digitorum longus muscles from WT and *Fam210a*^MKO^ mice were fixed in fixative containing 2.5% glutaraldehyde, 1.5% paraformaldehyde in 0.1 M sodium cacodylate buffer. Samples were rinsed in deionized water three times followed by 1% osmium tetroxide and 0.8% Ferricyanide solution in water to fix lipids and lipid-associated membranes for 2 hours. Then the samples were washed three times in water followed by 1% uranyl acetate in water for 20 minutes. After three washes in water, the samples were dehydrated in gradient ethanol solution from 50%, 75%, 95% to 100%. The samples were then washed three times in acetonitrile for transition and infiltration with acetonitrile:resin=2:1 mixture overnight. Then the samples were further infiltrated with acetonitrile:resin=1:2 for 3 hours before embedding in resin (Embed 812:DDSA:DMA=5:4:2, 0.22DMP-30). Utrathin sections were cut at 70 nm and counter stained with uranyl acetate and lead citrate. Stained sections were imaged under Tecnai T12 transmission electron microscope equipped with Gatan imaging system.

### Hematoxylin and Eosin and Immunofluorescence Staining

Muscle tissues were dissected and frozen immediately in O.C.T compound. Frozen samples were cross sectioned at 10 μm using Leica CM1850 cryostat. For hematoxylin and eosin staining, the slides were first stained with hematoxylin for 15min, rinsed in running tap water until clear and stained with eosin for 1min. Slides were dehydrated in gradient ethanol and then xylene before covered with Permount. For fiber type staining, the slides were directly incubated in blocking buffer for better signal detection. For other immunofluorescence staining, slides were fixed in 4% paraformaldehyde for 10minutes followed by 100 mM glycine in PBS to quench the excess PFA. Then slides were then incubated in blocking buffer (5% goat serum, 2% bovine serum albumin, 0.1% Triton X-100, and 0.1% sodium azide in PBS) for at least 1 hour at RT. Samples were incubated with indicated primary antibodies diluted in blocking buffer overnight at 4°C. After washing with PBST (0.1% Tween-20), samples were incubated with secondary antibodies and DAPI for 1hour at RT followed by PBST washes and cover in mounting media for imaging.

All H&E staining images were captured by a Nikon D90 digital camera mounted on a microscope with a 20X objective and all immunofluorescence images were captured using a Leica DM 6000B microscope with a 20X objective.

### Real-time PCR

Total RNA was extracted from muscle tissues using Trizol reagent according to the manufacturer’s instructions. 2 μg of total RNA was reverse transcribed using random primers with M-MLC reverse transcriptase (Invitrogen, cat#28025021). Real-time PCR was conducted on Roche Lightcycler 96 Real time PCR system (Roche) with FastStart Essential DNA Green Master (Roche, cat#06402712001). The 2^-ΔΔCt^ method was used to analyze the relative changes in each gene’s expression normalized to 18S expression.

### Protein Extraction and Western Blot Analysis

Total protein was isolated from tissues using RIPA buffer (50 mM Tris-HCl pH 8.0, 150 mM NaCl, 1% SDS, 1 mM EDTA, 0.5% NP-40, 0.5% sodium deoxycholate and 0.1% sodium dodecyl sulfate). Protein concentrations were determined by Pierce BCA Protein Assay Reagent (Thermo Scientific, cat# 23225). Proteins were resolved on SDS-PAGE, transferred onto 0.22 μm polyvinyldene fluoride (PVDF) membrane, blocked with 5% BSA for 1hour at RT, and then incubated with primary antibodies diluted in 5% BSA overnight at 4°C. Membranes were washed in TBST (0.1% Tween-20) followed by secondary antibody incubation for 1hour at RT. Immunodetection was performed using enhanced Western Blotting Chemiluminescence Luminol Reagent (Santa Cruz Biotechnology, cat#sc-2048) and detected with a FluorChem R System (ProteinSimple).

### Acetyl-CoA Measurement

Muscle lysate acetyl-CoA contents were measured by PicoProbe Acetyl-CoA fluorometric Assay Kit (BioVision Inc, cat#K317) according to the manufacturer’s instructions. In brief, muscle tissues from mice were weighed and snap frozen in liquid nitrogen immediately and then pulverized. Perchloric acid were used to homogenize the tissues. After centrifuge, KHCO_3_ were added to neutralize the supernatant. After brief centrifuge to pellet the KClO_4_, 10 μL of supernatant were added for the assay. Free CoASH, malonyl CoA and Succ-CoA in the samples were quenched by CoA quencher. The acetyl-CoA were then converted to CoA which reacted to form NADH which interacted with PicoProbe to generate fluorescence at Ex/Em=535/587 nm with Spark 10M multimode microplate reader (TECAN), and the acetyl-CoA concentration was calculated based on the standards following the manufacturer’s instructions.

### Seahorse Mitochondrial Respiration Analysis

Mitochondrial respiration was measured with isolated mitochondria from muscle in Seahorse XFe24 Analyzer (Agilent Technologies). The protocol was adapted from Agilent (Isolated Mitochondria Assay using the XF24 Analyzer). Briefly, mitochondria were isolated from muscles of the mouse and seeded at 5μg/well in 50 μL on the XFe24 cell culture microplate and centrifuged at 2,500 rpm for 20 minutes at 4°C. MAS-1 buffer (70 mM sucrose, 220 mM Mannitol, 5 mM KH_2_PO_4_, 5 mM MgCl_2_, 2 mM HEPES, 1 mM EGTA, 0.2% fatty acid-free BSA, pH 7.4) was used to prepare the buffer and reconstitute the drugs for the experiments. Then 450 μL of initial buffer (For electron flow experiment: 10 mM succinate, 2 μM Rotenone in MAS-1 buffer; for coupling experiment: 10 mM pyruvate, 2 mM malate, 4 μM FCCP in MAS-1 buffer) were added to each well. The mitochondria were then incubated for 8 minutes in a 37°C non-CO_2_ oven before the assay. Drug concentrations were as followed: for electron flow experiment: 20 μM Rotenone, 100 mM succinate, 40 μM Antimycin A, 100 mM Ascorbate+1 mM TMPD; for coupling experiment: 20 mM ATP, 20 μM oligomycin, 40 μM FCCP, 40 μM Antimycin A. Mitochondria respiration rates were generated by Seahorse Wave software.

### TCA cycle metabolomics

The TCA cycle metabolite detection protocol was adapted from ^56^. Briefly, muscle tissue was weighed and homogenized in Precellys CK14 tubes with 10volume of ice-cold 80% Methanol. Then the lysates were vortexed for 10 minutes. The homogenate was transferred to a new tube and an equal volume of ice-cold 80% methanol. Internal standard solutions were prepared as described in the protocol, with α-ketoglutarate, fumarate and succinate only. 10 μL of internal standard was added into the diluted homogenate. The homogenate was vortexed for 10mintes before clearing at 13,000 rpm for 10minutes. The resulting supernatant was dried with N_2_ for about 6 hours. The pellet was then reconstituted with 100 μL H_2_O. Then the samples were mixed with 50 μL 1M O-BHA (O-benzylhydroxylamine), 50 μL 1M EDC (1-ethyl-3-(3-dimethylaminopropyl)-carbodiimide) and vortexed for 1 hour at RT. Then 300 μL of ethyl acetate was added and vortexed for 10 minutes. The sample was then centrifuged at 13,000 rpm for 10 minutes and the top organic layer was collected and dried using a stream of N_2_ at 40°C and reconstituted in 100 μL of 50% methanol. The samples were then subjected to LC-MS detection.

### SCS activity assay

Succinyl-CoA synthetase activity was measured (Biovision, cat# K597-100) according to the manufacturer’s instructions. Briefly, mitochondria from muscles were isolated and used for experiments. 20 μg of mitochondria were resuspended and lysed in SCS assay buffer and used for experiments. The concentration of the resulting lysate was measured by BCA assay and an equal amount of protein were used for experiment.

### SDH staining

Muscle tissues were sectioned by cryosection and the slides were kept at -80°C until experiments. The slides were brought to RT from -80°C and incubated in incubation medium (270 mg sodium succinate, 10 mg nitro blue tetrazolium in 10 mL 0.2 M phosphate buffer, pH 7.6) for 60 minutes at 37°C. The slides were then washed with ddH_2_O for 3 times. Then the slides were incubated with increasing gradient of acetone 3 times from 30%, 60% to 90% acetone, each with 5minutes. When in last development of 90% acetone, leave the acetone until there was a faint purplish cloud. Then the slides were washed with decreasing gradient of acetone from 60% to 30%. Rinse the slides in waster, mount the slides and use for imaging.

### LC-MS/MS of muscle proteins and immuno-precipitated proteins

Mitochondria were purified from skeletal muscles and subjected to proteomics analysis. Or anti-acetylated-lysine antibody pull-down proteins from muscle lysate on agarose A/G beads were sent for LC-MS/MS to detect acetylated proteins. Briefly, the mitochondria pellet was lysed with 100 mM ammonium bicarbonate, supplemented with protease and phosphatase inhibitors. The protein concentration of the homogenate was determined by BCA assay. 500 μg of protein from each sample was precipitated with ice cold acetone at - 20°C overnight. The samples were then centrifuged at 12,000 g for 10 minutes. The pellets were reduced with 10 mM dithiothreitol prepared in 8 M urea, and alkylated with iodoethanol prepared in ACN buffer (2% iodoethanol, 0.5% triethylphosphine, 97.5% acetoniltrile). The proteins were digested with Trypsin/LysC mix (Promega, Wisconsin, WI, USA) at a 1:50 (w/w) enzyme-to-substrate ratio in a barocycler (Pressure BioScience) at 50°C with 60 cycles of 20 kpsi for 50 s and 1 atmospheric pressure for 10 seconds.

Samples were desalted using Pierce Peptide Desalting Spin Columns (Thermo Fisher Scientific). The samples were then dried and reconstituted in 20 μL of 0.1% formic acid (FA) in 3% acetonitrile. 1 μg of protein was loaded onto the column for proteomics analysis. The samples were analyzed by reverse-phase LC-ESI-MS/MS system using the Dionex UltiMate 3000 RSLC nano system, couple to a Q-Exactive High-Field (HF) Hybrid Quadrupole Orbitrap MS. The data was processed with MaxQuant Software. The mapped proteins were then compared to MitoCarta 3.0 ^57^ and only the mitochondrial proteins were considered for downstream analysis.

### Mitochondria Transfer Experiment

Mitochondria were isolated from the muscles and quantified according to the total protein amount using BCA reagent. Then the mitochondria were stained in MitoTracker Red (Cell Signaling Technology, cat#9082) for 30 min at 37°C for ease of visualizing the mitochondria, the mitochondria were then washed for two times before adding into the primary myoblast cell culture. Briefly 8μg of mitochondria were added into one well in a 24-well plate. The cells were collected at indicated time points and protein profile analyzed by western blot.

### Ribosome Isolation

Skeletal muscle from mice were dissected and flash frozen in LN_2_ until ready for experiment. The muscle was pulverized into powder and lysed in 3 volume of lysis buffer (0.25 M sucrose in medium N: 20 mM Tris-HCl pH 7.6, 100 mM KCl, 40 mM NaCl, 5 mM MgCl_2_, 6 mM β-mercaptoethanol). The lysates were passed through an 18-G syringe needle 10 times to ensure complete lysis. After 10 minutes incubation on ice, the lysates were cleared by centrifuge at 10,000 g for 15 min at 4°C. The supernatant was transferred and concentration measured by BCA assay. Equal concentrations between samples were adjusted using lysis buffer. The protein lysates were then treated with 1% Triton X-100 (final concentration) for 10 minutes on ice. 50 μL of the lysates were then aliquoted for input. The remaining lysates were loaded onto 1 M sucrose in medium N. The ribosomes were retrieved by centrifuging at 229,000 g for 20 hours at 4°C. The ribosomal pellet was dissolved in RIPA buffer and analyzed by SDS-PAGE.

### Western Blot SUnSET Experiment

Detailed protocol can be found in Craig A Goodman et al. 2013 ^34^. Briefly, mice were weighed and puromycin was injected at 0.04 μmol/g body mass. The mice were sacrificed immediately after 30 minutes and muscle samples were snap frozen in liquid nitrogen for western blot analysis. The proteins from muscle tissues were extracted with RIPA buffer and adjusted to the same concentration after BCA analysis. Proteins were analyzed by SDS-PAGE as previously mentioned.

### Polysome profiling experiment

Briefly, muscle tissues from mice were dissected and flash frozen in LN_2_ and kept at -80C until experiments. Muscles were pulverized with LN_2_ a pre-chilled mortar and pestle. The samples were lysed with hypotonic lysis buffer (5 mM Tris-HCl pH 7.5, 2.5 mM MgCl_2_, 1.5 mM KCl, 1000 μg/mL cycloheximide, 2 mM DTT, 0.5% Triton X-100, 0.5% Sodium Deoxycholate and protease inhibitor). The lysates were passed through an 18G syringe needle 10 times before sitting on ice for another 20 minutes to ensure complete lysis. Then the lysates were cleared of large debris by centrifuging at 10,000 g for 5 minutes at 4°C. The supernatants were collected and protein concentration measured by BCA assay. Equal amount and equal volume of proteins were loaded onto the 10%-50% sucrose gradient. The gradient was prepared by overlaying an equal volume 50% sucrose and 10% sucrose. The ultracentrifuge tubes were laid horizontal for 5 hours for the gradient to form ^58^. The proteins were separated at 36,000rpm for 16 hours at 4°C. The fractions were collected manually with 830 μL each. 10% TCA and 0.02% Sodium Deoxycholate were added to the fractions to precipitate the proteins O/N at 4°C. Then the pellet was spun down at 12,000 g for 10 minutes at 4°C. The protein pellets were washed with ice-cold acetone to get rid of the residual TCA. Pellets were then dried and dissolved in RIPA buffer. The proteins were analyzed on SDS-PAGE gel and the distribution of different subunits analyzed based on marker proteins.

### Statistical Analysis

Experiments involving mice were performed with a minimum of three biological replicates. All muscle histological analysis were quantified by Fiji-Image J software or by Adobe photoshop software. Skeletal muscle myofiber size quantification was done using Myosight ^59^. The myofiber size distribution graphs were made by joyplot in python. All experimental data were represented as mean ± SEM. Statistical significance was determined by the two tail Student *t* test. *p* value of less than 0.05 was considered as significant.

## Supporting information

Acetylated lysine antibody-IPMSMS

Fam210a-IPMS

Fam210aMKO muscle mitochondrial proteomics

primer table

## Acknowledgments

We thank Xiaoguang Zhu for assistance with confocal imaging, Jun Wu for mouse colony maintenance and management. We are grateful for Jill Hutchcroft at Purdue Flow Cytometry and Cell Separation, Amber Jannasch at Metabolite profiling, Uma Aryl at Bindley Mass Spectrometry Center, Laurie Mueller and Robert Seiler at Electron Microscopy and Imaging Facilities.

## Author contributions

Conceptualization: FY, SK

Methodology: JC, FY, KHK, PZ, JQ, WAT

Investigation: JC, KHK, FY, PZ, JQ

Visualization: JC, FY

Supervision: FY, SK

Writing—original draft: JC

Writing—review & editing: FY, SK

**Competing interests:** Authors declare that they have no competing interests.

## Data and materials availability

The datasets used for the analysis of *FAM210A* expression level are available in Gene Expression Omnibus (GEO): GDS473, GDS288, GDS2083, GDS4410, GDS611, GDS3637, GDS1879. No novel code was written for the analysis of the datasets used.

All data are available in the main text or the supplementary materials from the lead contact upon request.

## Supplementary figure legends

**Figure S1.**
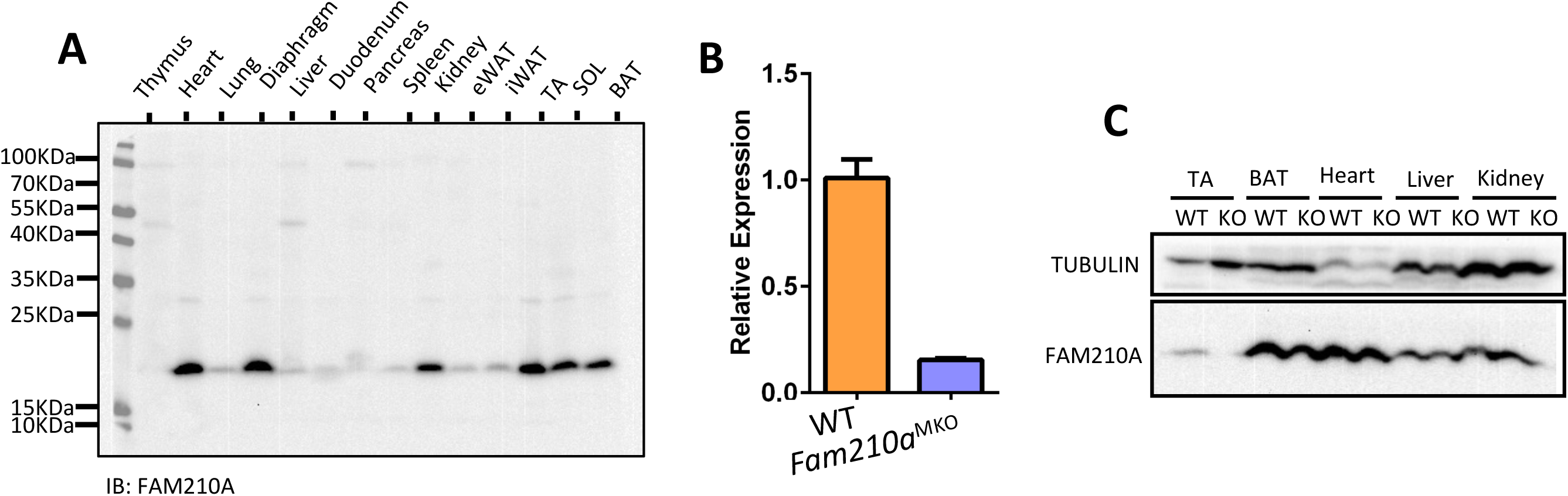
FAM210A tissue expression profile and validation of KO model. (**A**) Western blot analysis of FAM210A expression profile in different tissues in mice. (**B**) qPCR validation of the knockout of *Fam210a* skeletal muscle at 4 weeks of age. (**C**) Western blot validation of the knockout of FAM210A in skeletal muscle and other tissues.

**Figure S2.**
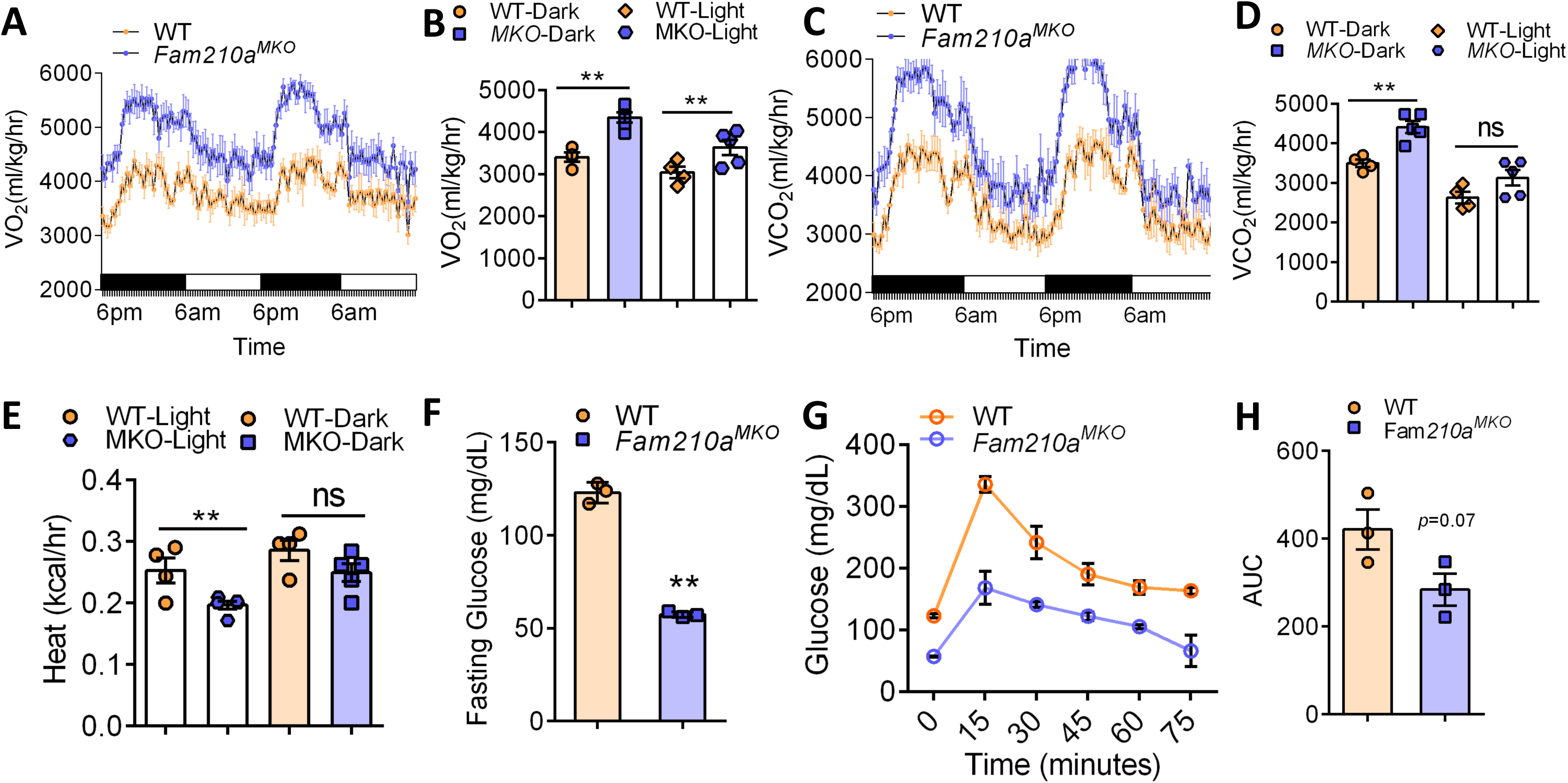
*Fam210a* loss results in systemic metabolic defects. (**A**) Metabolic chamber analysis of VO_2_ in WT and *Fam210a*^MKO^ mice at 4 weeks of age. (**B**) Bar graph of VO_2_ in WT and *Fam210a*^MKO^ mice, related to (**A**). (**C**) Metabolic chamber analysis of VCO_2_ in WT and *Fam210a*^MKO^ mice at 4 weeks of age. (**D**) Bar graph of VCO_2_ in WT and *Fam210a*^MKO^ mice related to (**C**). (**E**) Heat production of WT and *Fam210a*^MKO^ mice at 4 weeks of age. (**F**) Fasting glucose level in WT and *Fam210a*^MKO^ mice at 4 weeks of age. (**G**) Glucose tolerance test in WT and *Fam210a*^MKO^ mice at 4 weeks of age. (**H**) Area under curve analysis of GTT, related to (**G**).

**Figure S3.**
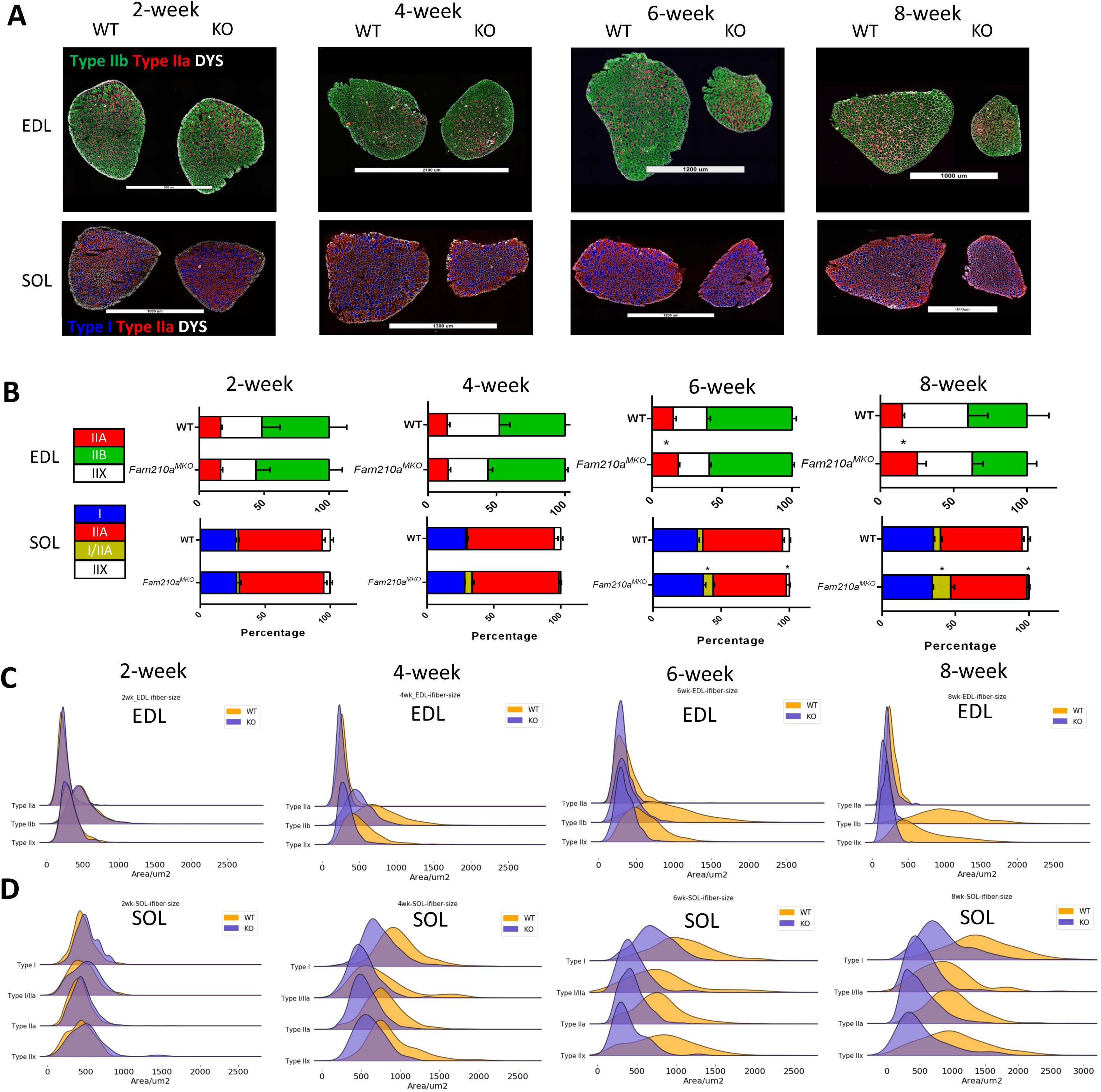
Fiber type analysis of the fast and slow muscles of Fam210aMKO mice at different ages. (**A**) Fiber type staining of EDL and SOL muscle at different ages. (**B**) Quantification of the composition of fiber type at different ages related to (A). (**C**) Fiber type specific fiber size distribution in EDL muscle in WT and *Fam210a*^MKO^ mice at different stages. (**D**) Fiber type specific fiber size distribution in SOL muscle in WT and *Fam210a*^MKO^ mice at different stages.

**Figure S4.**
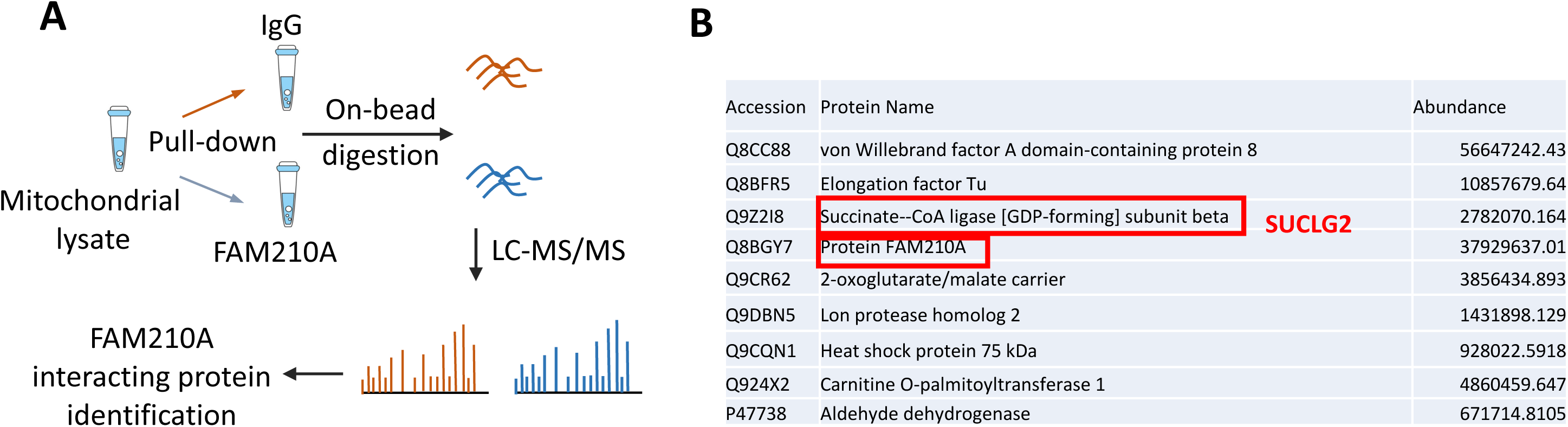
Identification of FAM210A interacting proteins. (**A**) Workflow of FAM210A antibody pull-down to identify FAM210A interacting proteins by LC-MS/MS. (**B**) Potential candidates for FAM210A interacting proteins.

**Figure S5.**
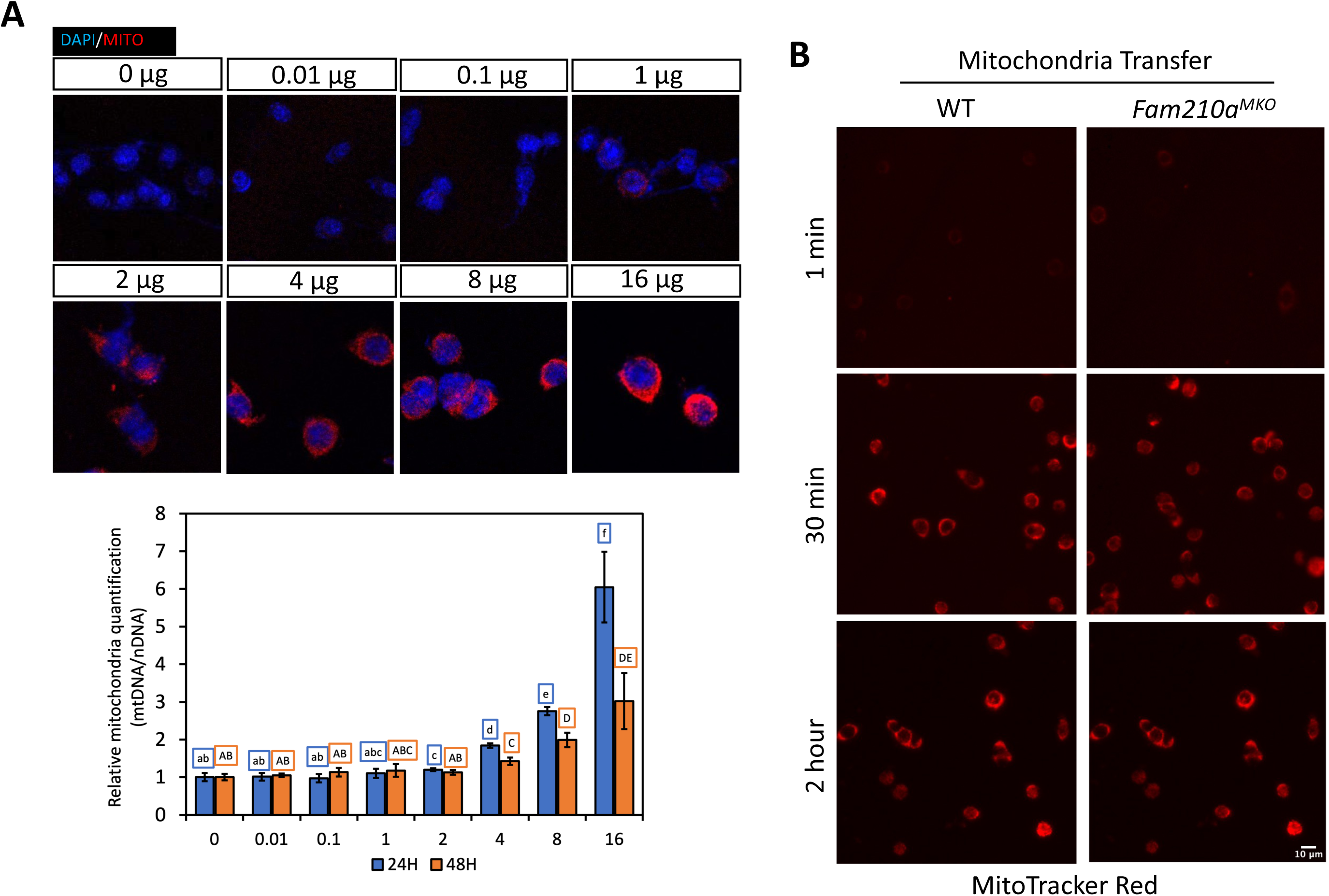
Mitochondria readily enter recipient cells after transplantation. (**A**) Different quantities of the MitoTrackerRed stained mitochondria after transplant into WT recipient myoblasts. Quantification of the mtDNA copy number relative to the nDNA after different lengths of transplantation shown in the bar graph. (**B**) MitoTrackerRed staining in WT recipient myoblasts at different timepoints after the transfer of mitochondria from WT and *Fam210a*^MKO^ muscle.

