## Supplementary material for "FAM210A mediates an inter-organelle crosstalk essential for protein synthesis and muscle growth in mouse": primer table

| Primer | Sequence (5’-3’) |
| --- | --- |
| q-cGAS-F | GTTCAAACACAAGAAATGCACTG |
| q-cGAS-R | GCTGACGGAGTACACAATCCT |
| q-Sting-F | TGAAAGGCTCTTCATTGTCTCTT |
| q-Sting-R | TGGCATCTTCTGCTTCCTAGA |
| q-Pink1-F | TTCTTCCGCCAGTCGGTAG |
| q-Pink1-R | CTGCTTCTCCTCGATCAGCC |
| q-Perk-F | GCGTCGGAGACAGTGTTTG |
| q-Perk-R | CGTCCATCTAAAGTGCTGATGAT |
| q-sXbp1-F | ACACGTTTGGGAATGGACAC |
| q-sXbp1-R | CCATGGGAAGATGTTCTGGG |
| q-Tafazzin-F | CCCCCGCTTTGGACAGAAAAT |
| q-Tafazzin-R | AGGCTGGAAATGATTGTGGAG |
| q-Fam210a-F | TGACAGCCTACGCCATGTTT |
| q-Fam210a-R | GGGTTGACATGTAGCCGTGA |
| q-Sirt5-F | CTCCGGGCCGATTCATTTCC |
| q-Sirt5-R | GCGTTCGCAAAACACTTCCG |
| q-Alas1-F | TCGCCGATGCCCATTCTTATC |
| q-Alas1-R | GGCCCCAACTTCCATCATCT |
| q-Tbk1-F | ACTGGTGATCTCTATGCTGTCA |
| q-Tbk1-R | TTCTGGAAGTCCATACGCATTG |
| q-Fgf21-F | TGACGACCAAGACACTGAAGC |
| q-Fgf21-R | TTTGAGCTCCAGGAGACTTTCTG |
| q-Atf4-F | AGCAAAACAAGACAGCAGCC |
| q-Atf4-R | ACTCTCTTCTTCCCCCTTGC |
| q-mtDNA-Nd2-F | CCTATCACCCTTGCCATCAT |
| q-mtDNA-Nd2-R | GAGGCTGTTGCTTGTGTGAC |
| q-nDNA-Pecam-F | ATGGAAAGCCTGCCATCATG |
| q-nDNA-Pecam-R | TCCTTGTTGTTCAGCATCAC |
| q-BiP(Hspa5)-F | TCATCGGACGCACTTGGAA |
| q-BiP(Hspa5)-R | CAACCACCTTGAATGGCAAGA |
